# Single Cell Transcriptional Signatures of the Human Placenta in Term and Preterm Parturition

**DOI:** 10.1101/738658

**Authors:** Roger Pique-Regi, Roberto Romero, Adi L.Tarca, Edward D. Sendler, Yi Xu, Valeria Garcia-Flores, Yaozhu Leng, Francesca Luca, Sonia S. Hassan, Nardhy Gomez-Lopez

## Abstract

More than 135 million births occur each year; yet, the molecular underpinnings of human parturition in gestational tissues, and in particular the placenta, are still poorly understood. The placenta is a complex heterogeneous organ including cells of both maternal and fetal origin, and insults that disrupt the maternal-fetal dialogue could result in adverse pregnancy outcomes such as preterm birth. There is limited knowledge of the cell type composition and transcriptional activity of the placenta and its compartments during physiologic and pathologic parturition. To fill this knowledge gap, we used scRNA-seq to profile the placental villous tree, basal plate, and chorioamniotic membranes of women with or without labor at term and those with preterm labor. Significant differences in cell type composition and transcriptional profiles were found among placental compartments and across study groups. For the first time, two cell types were identified: 1) lymphatic endothelial decidual cells in the chorioamniotic membranes, and 2) non-proliferative interstitial cytotrophoblasts in the placental villi. Maternal macrophages from the chorioamniotic membranes displayed the largest differences in gene expression (e.g. NFKB1) in both processes of labor; yet, specific gene expression changes were also detected in preterm labor. Importantly, several placental scRNA-seq transcriptional signatures were modulated with advancing gestation in the maternal circulation, and specific immune cell type signatures were increased with labor at term (NK-cell and activated T-cell) and with preterm labor (macrophage, monocyte, and activated T-cell). Herein, we provide a catalogue of cell types and transcriptional profiles in the human placenta, shedding light on the molecular underpinnings and non-invasive prediction of the physiologic and pathologic parturition.

**One sentence summary:** The common molecular pathway of parturition for both term and preterm spontaneous labor is characterized using single cell gene expression analysis of the human placenta.

## Main text

Parturition is essential for the reproductive success of viviparous species^1^; yet, the mechanisms responsible for the onset of labor remain to be elucidated^2, 3^. Understanding human parturition is essential to tackle the challenge of prematurity, which affects 15 million neonates every year^4–6^. Bulk transcriptomic studies of the cervix^7–11^, myometrium^12–17^, and chorioamniotic membranes^18–20^ revealed that labor is a state of physiological inflammation; however, finding specific pathways implicated in preterm labor still remains an elusive goal. A possible explanation is that gestational tissues, and especially the placenta, are heterogeneous composites of multiple cell types, and elucidating perturbations in the maternal-fetal dialogue requires dissection of the transcriptional activity at the cell type level, which is not possible using bulk analyses. Recent microfluidic and droplet-based technological advances have enabled characterization of gene expression at single-cell resolution (scRNA-seq)^21, 22^. Previous work in humans^23–25^ and mice^26^ demonstrated that scRNA-seq can capture the multiple cell types that constitute the placenta and identify their maternal or fetal origin. Such studies showed that single-cell technology can be used to infer communication networks across the different cell types at the maternal-fetal interface^25^. Further, the single-cell-derived placental signatures were detected in the cell-free RNA present in maternal circulation^23^, suggesting that non-invasive identification of women with early-onset preeclampsia is feasible. However, these studies included a limited number of samples and did not account for the fact that different pathologies can arise from dysfunction in different placental compartments.

In addition, the physiologic and pathologic processes of labor have never been studied at single-cell resolution.

In this study, a total of 25 scRNA-seq libraries were prepared from three placental compartments: basal plate (BP), placental villous (PV), and chorioamniotic membranes (CAM) (Figure 1A). These were collected from 9 women in the following study groups: term no labor (TNL), term in labor (TIL), and preterm labor (PTL). scRNA-seq libraries were prepared with the 10X Chromium system and were processed using the 10X Cell Ranger software, resulting in 79,906 cells being captured and profiled across all samples (Table S1). We used Seurat^27^ to normalize expression profiles and identified 19 distinct clusters, which were assigned to cell types based on the expression of previously reported marker genes^23–25^ (see Methods, Figure S1 and Table S2-3). The uniform manifold approximation and projection (UMAP^28^) was used to display these clusters in two dimensions (Figure 1B). With this approach, the local and global topological structure of the clusters is preserved, with subtypes of the major cell lineages (trophoblast, lymphoid, myeloid, stromal, and endothelial sub-clusters) being displayed proximal to each other. The trophoblast lineage shown in Figure 1B recapitulates the differentiation structure previously reported^23, 25^, such as the progression from cytotrophoblasts to either extravillous trophoblasts or syncytiotrophoblasts (Figure S2).

**Figure 1.**
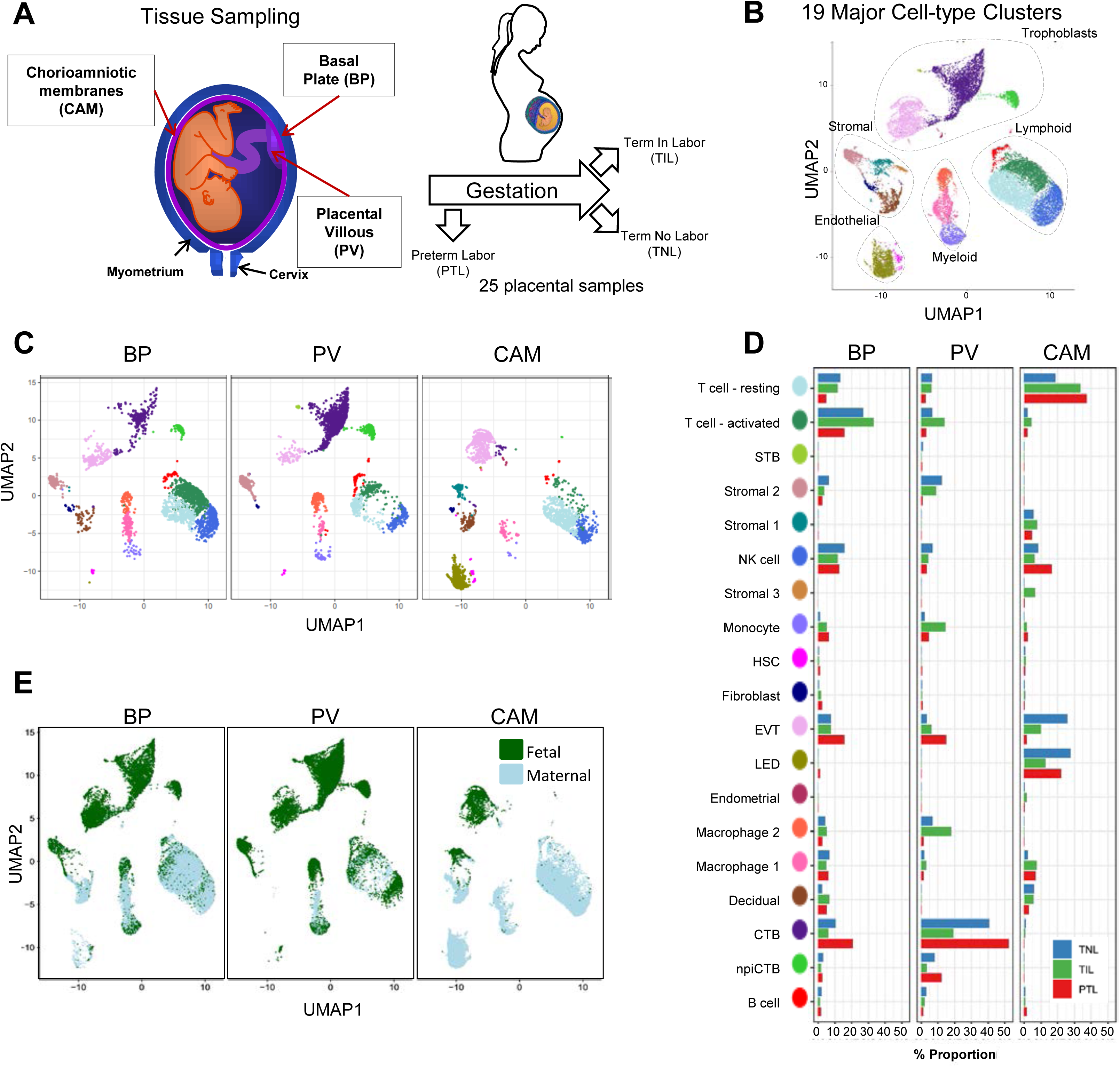
Transcriptional map of the placenta in human parturition. **A**. Study design illustrating the placental compartments and study groups. **B**. Uniform Manifold Approximation Plot (UMAP), where dots representing single-cells and colored by cell type. **C**. Distribution of single-cell clusters by placental compartments. **D**. Average proportions of cell types by placental compartments and study groups. **E**. Distribution of single-cells by maternal or fetal origin. STB, Syncytiotrophoblast; EVT, Extravillous trophoblast; CTB, cytotrophoblast; HSC, hematopoietic stem cell; npiCTB, non proliferative interstitial cytotrophoblast; LED, lymphoid endothelial decidual cell

The magnitude of cell type composition differences across placental compartments was substantial (Figure 1C); yet, significant changes among study groups were also noted (Figure 1D). Extravillous trophoblasts (EVT) were present in all three compartments, while cytotrophoblasts (CTB) were especially pervasive in the placental villi. Non-proliferative interstitial cytotrophoblasts (npiCTB) were identified for the first time in the placental villi as forming a distinct cluster. Compared to conventional cytotrophoblasts, npiCTBs have a higher expression of PAGE4, but a reduced expression of genes involved in cell proliferation such as XIST, DDX3X, and EIF1AX (Figure S3).

In terms of immune cell types, the chorioamniotic membranes largely contained lymphoid and myeloid cells of maternal origin, including T cells (mostly in a resting state), NK cells, and macrophages (Figure 1C, 1E and Figure S4). In contrast, the basal plate included immune cells of both maternal and fetal origin, such as T cells (mostly in an active state), NK cells, and macrophages. The placental villi contained more fetal than maternal immune cells, namely monocytes, macrophages, T cells, and NK cells. Two macrophage subsets were found in placenta compartments: macrophage 1 of maternal origin that was predominant in the chorioamniotic membranes, and macrophage 2 of fetal origin that was mainly present in the basal plate and placental villi.

Importantly, a new lymphatic endothelial decidual (LED) cell type of maternal origin was identified in the chorioamniotic membranes, forming a distinct transcriptional cluster that was separate from other endothelial cell-types (Figure 1C). LED cells were rarely observed in the basal plate and were completely absent in the placental villous (Figure 1D). The signature genes of this novel cell type were enriched for pathways involving cell surface interactions at the vascular wall, extracellular matrix organization (Figure S5), tight junction, and focal adhesion (Figure S6). Immunostaining confirmed the co-expression of LYVE1 (lymphatic marker) and CD31 (endothelial molecule marker) in the vessels of the chorioamniotic membranes, but not in the basal plate or placenta (Figure 2A). The co-expression of LYVE1 and CD31 (i.e. LED cells) in the chorioamniotic membranes is shown in Figure 2B and Video S1. LYVE1 was also expressed by the fetal macrophages present in the placental villi and basal plate (Figure 2C), yet was only visualized by immunostaining in immune cells located in the villous tree (Figure 2A). Other genes highly expressed by LED cells were CD34, CDH5, EDNRB, PDPN, and TIE1 (Figure 2C, S7). This finding conclusively shows the presence of lymphatic vessels in the decidua parietalis of the chorioamniotic membranes, providing a major route for maternal T cells infiltrating the maternal-fetal interface^29^.

**Figure 2.**
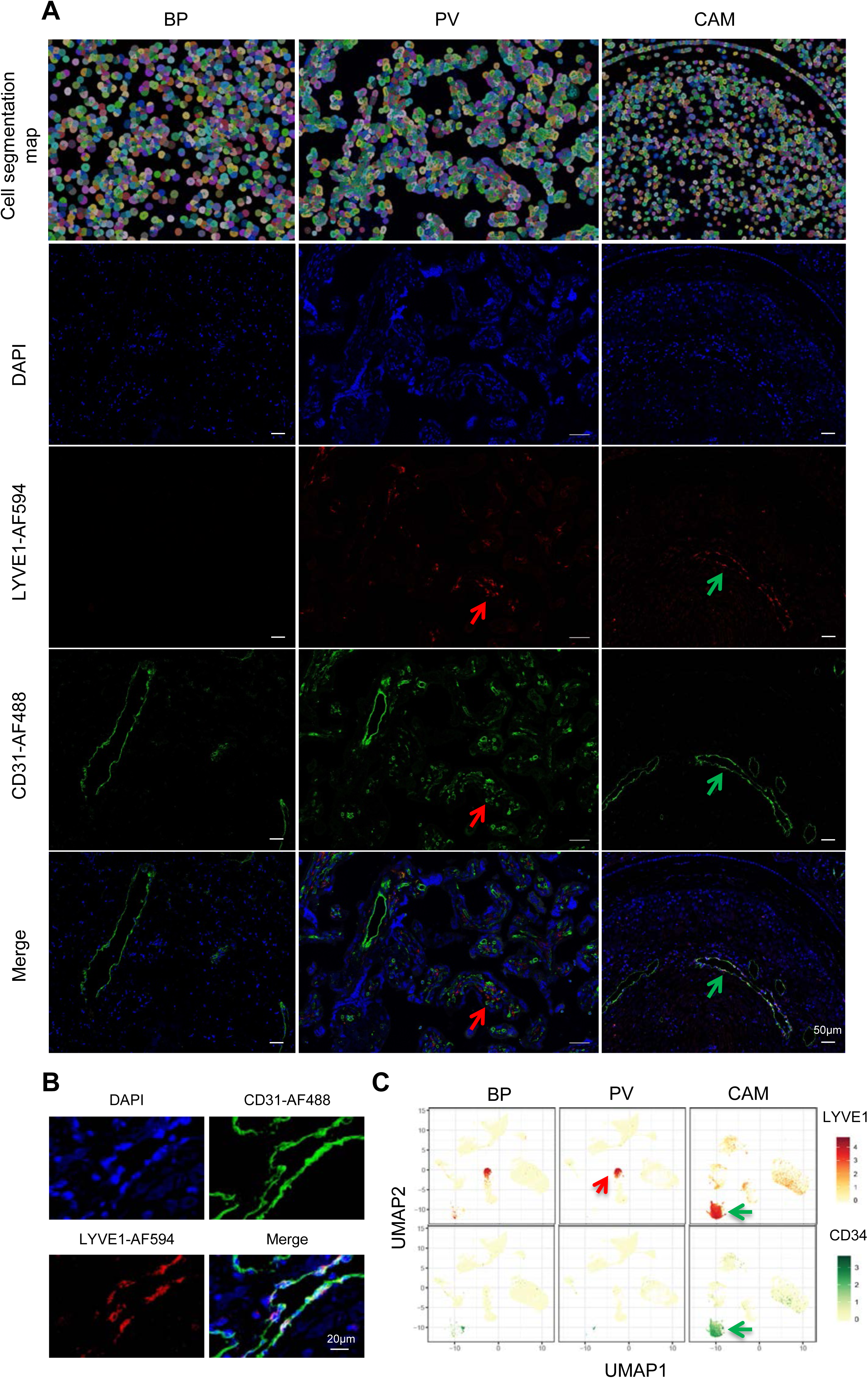
Identification of LED cells in the chorioamniotic membranes. **A**. Cell segmentation map (built using the DAPI nuclear staining) and immunofluorescence detection of LYVE-1 (red) and CD31 (green) in the basal plate (BP), placental villi (PV), and chorioamniotic membranes (CAM). Red arrows point to fetal macrophages expressing LYVE1 but not CD31, and green arrows indicate lymphatic endothelial decidual cells expressing both LYVE1 and CD31. **B**. Co-expression of LYVE1 and CD31 (i.e. LED cells) in the chorioamniotic membranes. **C**. Single-cell expression UMAP of LYVE-1 (red) and CD34 (green) in the placental compartments.

For cell types that were present in more than one placental compartment, major differences in gene expression were identified across locations, indicative of further specialization of cells depending on the unique physiological functions of each microenvironment (Figure S8 and Table S4). Differences in the transcriptional profiles were particularly large for maternal macrophages as well as EVTs, NK cells, and T cells in the chorioamniotic membranes compared to the other compartments. Genes differentially expressed in the chorioamniotic membranes were enriched for interleukin, Toll-like receptor, and the NF-κB and TNF signaling pathways (Figure S9-S11). These results are consistent with previous reports showing a role for these mediators in the inflammatory process of labor^30–51^. Conversely, the placental villous and basal plate were more similar to each other, with most differentially expressed genes (DEG) between these compartments being noted in fibroblasts (335 DEG, q<0.1 and fold change >2) (Figure S8, S12-S17). DEGs in the placental villous fibroblasts showed enrichment in smooth muscle contraction, apelin and oxytocin signaling pathways (Figure S16); while DEGs in CAM fibroblasts were enriched in elastic fiber formation and extracellular matrix pathways (Figure S9).

Next, we assessed changes due to term and preterm labor in each cell type (Table S5). The largest number of DEGs between the term labor and term no labor groups were observed in the maternal macrophages (macrophage 1), followed by the EVT (144 and 37, respectively, q<0.1; Figure 3A). The largest number of DEGs between the preterm labor and term labor groups were observed in EVT and CTB (37 and 33, respectively, q<0.1; Figure 3A). Figure 3B displays the gene expression changes between TIL and TNL or PTL and TNL that are shared between the two labor groups, representing the common pathway of parturition (defined as the anatomical, physiological, biochemical, endocrinological, immunological, and clinical events that occur in the mother and/or fetus in both term and preterm labor^52^). Non-shared differences in gene expression with labor at term and in preterm labor were mostly observed in trophoblast cell types such as CTB and EVT as well as in stromal cells (Figure 3C). Some of these changes may be explained by the unavoidable confounding effect of gestational age since placentas from women without labor in preterm gestation cannot be obtained in the absence of pregnancy complications. Specifically, the expression of NFKB1 by maternal macrophages was higher in labor at term compared to TNL, and this increase was further accentuated in preterm labor (Figure 3D). Consistently with the induction of the NFκB pathway, the labor-associated DEGs in macrophages involved biological processes such as activation of immune response and regulation of cytokine production (Figure S18A). When comparing the effect sizes between the PTL/TNL and TIL/TNL juxtapositions on the same gene and cell type, positive correlations were observed for most of the placental cell types (Figure 3E). Genes displaying differential effects in term and preterm labor are mostly found in trophoblast cell types (see off-diagonal points in the scatter plot), which may be explained by the phenomenon of gene expression decoherence^53^. This lack of proper correlation between biomarkers to their expected normal relationships is commonly found in pathological conditions.

**Figure 3.**
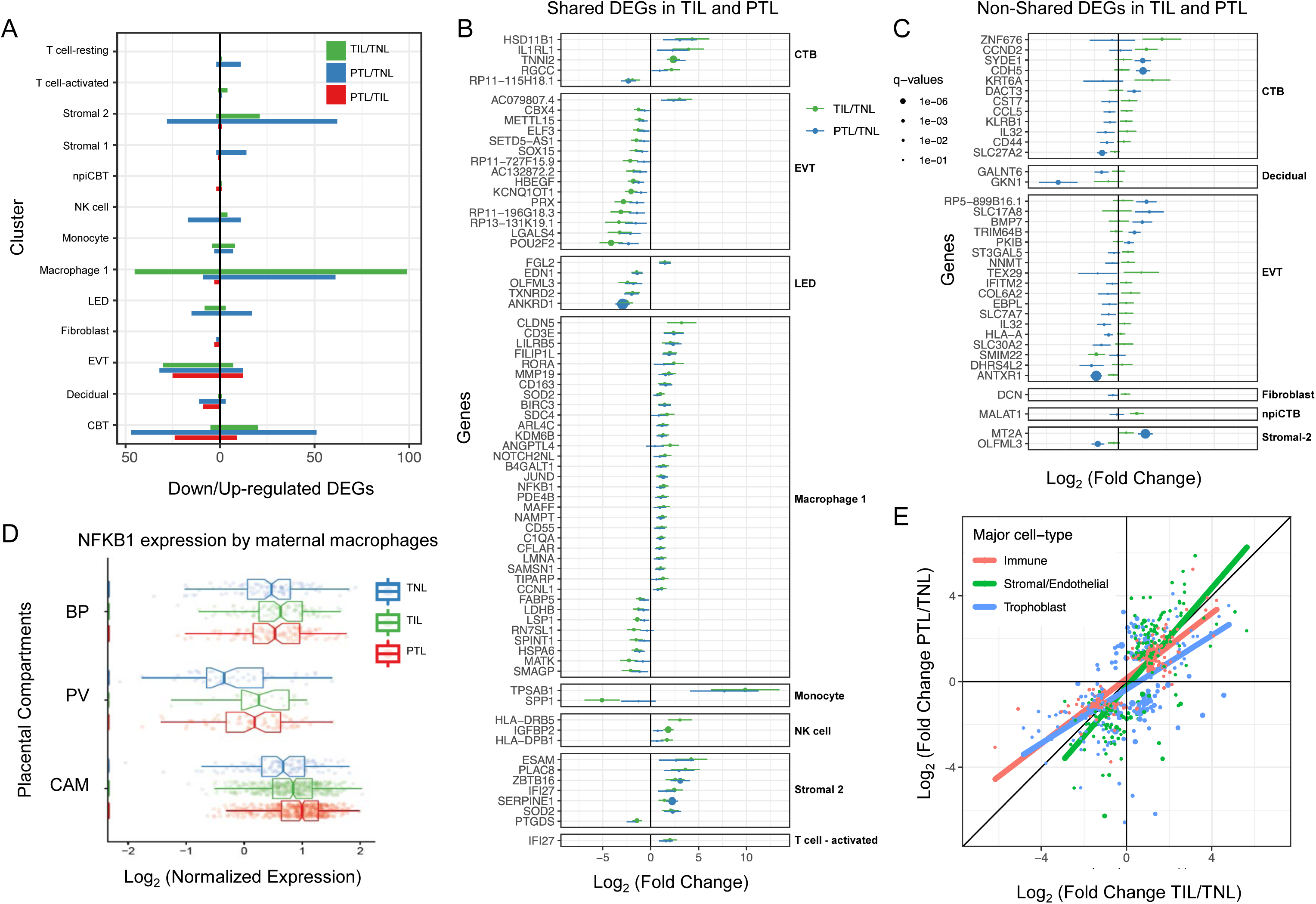
Cell type specific expression changes in term and preterm labor. **A**. Number of differentially expressed genes (DEGs) among study groups (TNL, term no labor; TIL, term in labor; PTL, preterm labor) by direction of change. Shared (**B**) and non-shared (**C**) expression changes in term labor and preterm labor relative to the term no labor group (q<0.01). The length of each whisker represents the 95% confidence interval. **D**. The expression of NFKB1 by maternal macrophages in the placental compartments (BP, basal plate; PV, placental villous; CAM, chorioamniotic membranes) and study groups. The notch represents the 95% confidence interval of the median. **E**. Differences and similarities in expression changes with preterm labor and term labor by three major cell types (immune, stromal/endothelial, and trophoblast cells).

Lastly, in EVT the DEGs with labor were enriched for genes implicated in cellular response to stress, including the WNT and NOTCH pathways, as well as cell cycle checkpoints (Figure S18B), further supporting the hypothesis that the cellular senescence pathway (i.e. cell cycle arrest) implicated in the physiologic^54, 55^ and pathologic^56, 57^ processes of labor.

To demonstrate the translational value of single-cell RNA signatures derived from the placenta, we conducted an *in silico* analysis in public datasets^58, 59^ to test whether the single-cell signatures could be non-invasively monitored in the maternal circulation throughout gestation (Figure 4A). Previous studies have correlated bulk mRNA expression in the maternal circulation with gestational age at blood draw^58, 60^, risk for preterm birth^59, 61–63^, or both^64, 65^. First, we showed that the single-cell signatures of macrophages, monocytes, NK cells, T cells, npiCTB, and fibroblasts are modulated throughout gestation in the maternal circulation (Figure 4B-C, S19A). These results validate the T-cell and monocyte signature changes with gestational age that were previously reported^23, 58^; yet, here we show that novel placental single-cell signatures (e.g., npiCTB and fibroblast) can also be non-invasively monitored in maternal circulation (Figure S19A). In addition, for the first time, we report that the single-cell-derived NK-cell and activated T-cell signatures were upregulated in women with spontaneous labor at term compared to gestational-age matched controls without labor (Figure 4D). Importantly, at 24-34 weeks of gestation, we found that the single cell signatures of macrophages, monocytes, activated T cells, and fibroblasts were increased in the circulation of women with preterm labor and delivery compared to gestational age-matched controls (Figure 4E and S19B). These findings are in line with previous reports indicating a role for these immune cell-types in the pathophysiology of preterm labor^29, 66–68^.

**Figure 4.**
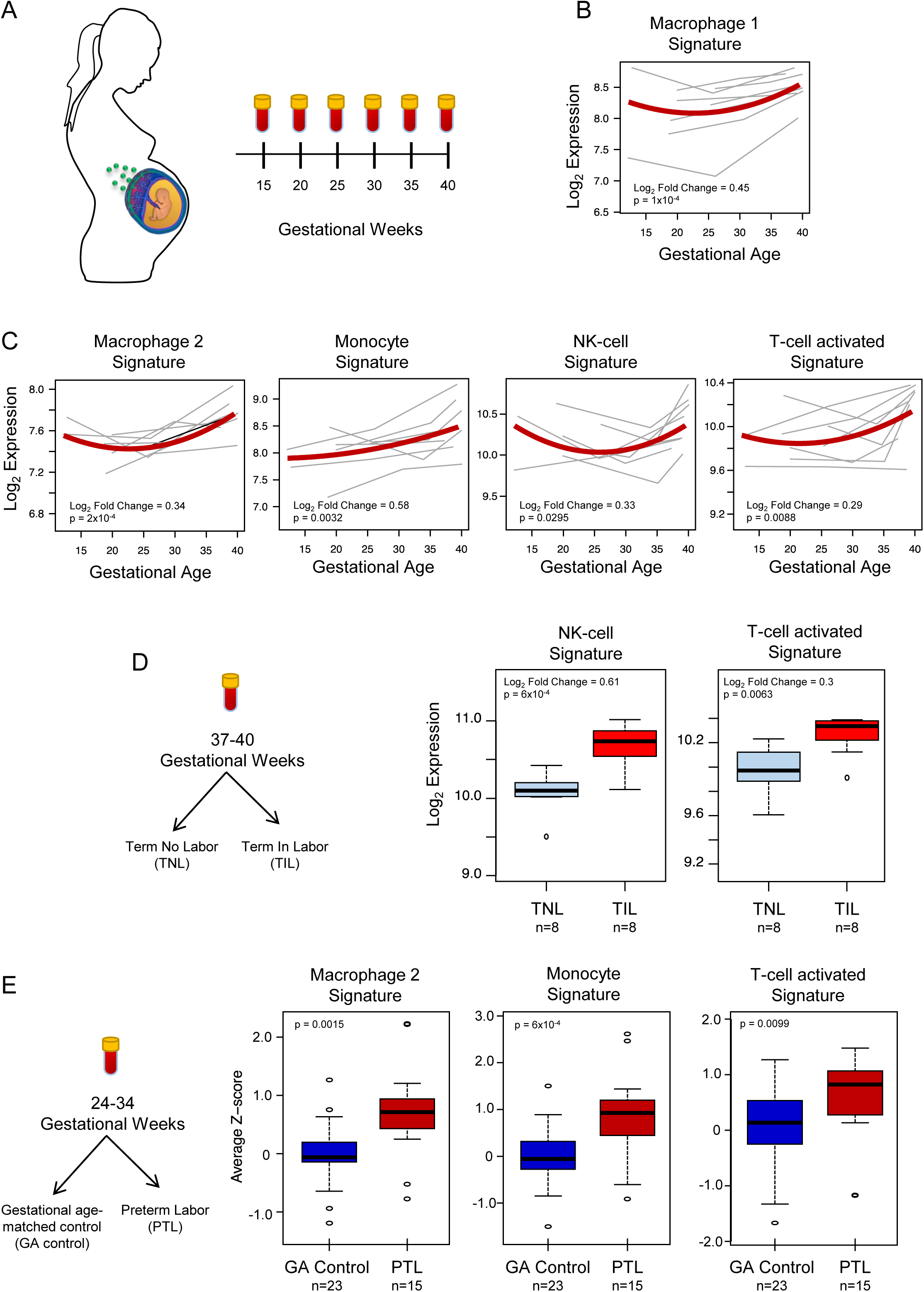
Quantification of scRNA-seq signatures in maternal circulation. A. Design of *in silico* longitudinal study including samples throughout gestation in women who delivered at term^58^. **B&C**. Variation of scRNA-seq signatures in the maternal circulation with advancing gestation. **D&E**. Perturbations in scRNA-seq signatures in women with spontaneous labor at term (TIL)^58^ or preterm labor (PTL)^59^. Gestational age-matched controls were included in each case (TNL, term no labor) and (GA control).

In summary, this study provides evidence of differences in cell type composition and transcriptional profiles among the basal plate, placental villi, and chorioamniotic membranes, as well as between the pathologic and physiologic processes of labor at single-cell resolution. Using scRNAseq technology, two novel cell types were identified in the chorioamniotic membranes and placental villi. In addition, we showed that maternal macrophages and extravillous trophoblasts are the cell types with the most transcriptional changes during the processes of labor. Lastly, we report that maternal and fetal transcriptional signatures derived from placental scRNA-seq are modulated with advancing gestation and are markedly perturbed with term and preterm labor in the maternal circulation. These results highlight the potential of single-cell signatures as biomarkers to non-invasively monitor the cellular dynamics during pregnancy and to predict obstetrical disease. The current study represents the most comprehensive single-cell analysis of the human placental transcriptome in physiologic and pathologic parturition; yet, additional studies are needed to characterize the different etiologies of the preterm labor syndrome.

## Data and materials availability

The scRNA-seq data reported in this study is being submitted to NIH dbGAP repository (accession number phsXXXXXX pending). All other data used in this study are already available through Gene Expression Omnibus (accession identifiers GSE114037 and GSE96083). All software and R packages used herein are detailed in the Materials and Methods. Scripts detailing the analyses are also available at https://github.com/piquelab/sclabor. To enable further exploration of the results we have also provided a Shiny App in Rstudio available at: http://genome.grid.wayne.edu:9091/

## Supplementary Materials

Materials and Methods

Tables S1-S6

Video S1

Figures S1-S19

## Materials and Methods

### Sample collection and processing, single-cell preparation, library preparation, and sequencing

#### Human subjects

Immediately after delivery, placental samples [the villi, basal plate (including the decidua basalis) and chorioamniotic membranes (including the decidua parietalis)] were collected from women with or without labor at term or preterm labor at the Detroit Medical Center, Wayne State University School of Medicine (Detroit, MI). Labor was defined by the presence of regular uterine contractions at a frequency of at least two contractions every 10 min with cervical changes resulting in delivery. Women with preterm labor delivered between 33-35 weeks of gestation whereas those with term labor delivered between 38-40 weeks of gestation (Table S6). The collection and use of human materials for research purposes were approved by the Institutional Review Boards of the Wayne State University School of Medicine. All participating women provided written informed consent prior to sample collection.

#### Single-cell preparation

Cells from placental villi, basal plate, and chorioamniotic membranes were isolated by enzymatic digestion, using previously described protocols with modifications^23, 69^. Briefly, placental tissues were homogenized using a gentleMACS Dissociator (Miltenyi Biotec, San Diego, CA) either in an enzyme cocktail from the Umbilical Cord Dissociation Kit (Miltenyi Biotec) or in collagenase A (Sigma Aldrich, St. Louis, MO). After digestion, homogenized tissues were washed with ice-cold 1X phosphate-buffered saline (PBS) and filtered through a cell strainer (Fisher Scientific, Durham, NC). Cell suspensions were then collected and centrifuged at 300 × g for 5 min. at 4°C. Red blood cells were lysed using a lysing buffer (Life Technologies, Grand Island, NY). Next, cells were washed with ice-cold 1X PBS and resuspended in 1X PBS for cell counting, which was performed using an automatic cell counter (Cellometer Auto 2000; Nexcelom Bioscience, Lawrence, MA). Lastly, dead cells were removed from the cell suspensions using the Dead Cell Removal Kit (Miltenyi Biotec) and cells were counted again using an automatic cell counter.

#### Single-cell preparation using the 10x Genomics platform

Viable cells were used for single-cell RNAseq library construction using the Chromium™ Controller and Chromium™ Single Cell 3’ version 2 kit (10x Genomics, Pleasanton, CA), following the manufacturer’s instructions. Briefly, viable cell suspensions were loaded into the Chromium™ Controller to generate gel beads in emulsion (GEM) with each GEM containing a single cell as well as barcoded oligonucleotides. Next, the GEMs were placed in the Veriti 96-well Thermal Cycler (Thermo Fisher Scentific, Wilmington, DE) and reverse transcription was performed in each GEM (GEM-RT). After the reaction, the complementary cDNA was cleaned using Silane DynaBeads (Thermo Fisher Scentific) and the SPRIselect Reagent kit (Beckman Coulter, Indianapolis, IN). Next, the cDNAs were amplified using the Veriti 96-well Thermal Cycler and cleaned using the SPRIselect Reagent kit. Indexed sequencing libraries were then constructed using the Chromium™ Single Cell 3’ version 2 kit, following the manufacturer’s instructions.

#### Library preparation

cDNA was fragmented, end-repaired, and A-tailed using the Chromium™ Single Cell 3’ version 2 kit, following the manufacturer’s instructions. Next, adaptor ligation was performed using the Chromium™ Single Cell 3’ version 2 kit followed by post-ligation cleanup using the SPRIselect Reagent kit to obtain the final library constructs, which were then amplified using PCR. After performing a post-sample index double-sided size selection using the SPRIselect Reagent kit, the quality and quantity of the DNA were analyzed using the Agilent Bioanalyzer High Sensitivity chip (Agilent Technologies, Wilmington, DE). The Kapa DNA Quantification Kit for Illumina® platforms (Kapa Biosystems, Wilmington, MA) was used to quantify the DNA libraries, following the manufacturer’s instructions.

#### Sequencing

Sequencing of the single-cell libraries was performed by NovoGene (Sacramento, CA) using the Illumina Platform (HiSeq X Ten System).

### Immunofluorescence

Samples of chorioamniotic membranes, placenta villi, and decidua basal plate were embedded in Tissue-Tek Optimum Cutting Temperature (OCT) compound (Miles, Elkhart, IN) and snap-frozen in liquid nitrogen. Ten-µm-thick sections of each OCT-embedded tissue were cut using the Leica CM1950 (Leica Biosystems, Buffalo Grove, IL). Frozen slides were thawed to room temperature, fixed with 4% paraformaldehyde (Electron Microscopy Sciences, Hatfield, PA), and washed with 1X PBS. Non-specific background signals were blocked using Image-iT FX Signal Enhancer (Life Technologies) followed by blocking with antibody diluent/blocker (Perkin Elmer, Waltham, MA) for 30 min. at room temperature. Slides were then incubated with the rabbit anti-LYVE-1 antibody (Novus Biologicals, Centennial, CO) and the Flex mouse anti-human CD31 antibody (clone JC70A, Dako North America, Carpinteria, CA) for 90 min. at room temperature. Following washing with 1X PBS and blocking with 10% goat serum (SeraCare, Milford, MA), the slides were incubated with secondary goat anti-rabbit IgG– Alexa Fluor 594 (Life Technologies) and goat anti-mouse IgG–Alexa Fluor 488 (Life Technologies) for 30 min. at room temperature. Finally, the slides were washed and coverslips were mounted using ProLong Gold Antifade Mountant with DAPI (Life Technologies). Immunofluorescence was visualized using a confocal fluorescence microscope (Zeiss LSM 780; Carl Zeiss Microscopy GmbH, Jena, Germany) at the Microscopy, Imaging, and Cytometry Resources Core at the Wayne State University School of Medicine. Tile scans were performed from the chorioamniotic membranes, placental villi, and basal plate and the complete imaging fields were divided into six-by-six quadrants.

### scRNA-seq data analyses

Raw fastq files obtained from Novogene were processed using Cell Ranger version 2.1.1 from 10X Genomics. First, sequence reads for each library (sample) were aligned to the hg19 reference genome using the STAR^70^ aligner, and expression of gene transcripts documented in the ENSEMBL database (Build 82) were determined for each cell. Gene expression was determined by the number of unique molecular identifiers (UMI) observed per gene. Second, data were aggregated and down-sampled to take into account differences in sequencing depth across libraries using Cell Ranger Aggregate to obtain gene by cell expression data. Third, Seurat^27^ was used to further clean and normalize the data. Then, only data from cells with a minimum of 200 detected genes, and from genes expressed in at least 10 cells were retained. Cells expressing mitochondrial genes at a level of >10% of total gene counts were also excluded, resulting in 77,906 cells and 25,803 genes (summary in Table S1). Gene read counts were normalized with the Seurat “NormalizeData” function (normalization.method = LogNormalize, scale.factor = 10,000). Genes showing significant variation across cells were selected based on “LogVMR” dispersion function and “FindVariableGenes”. Ribosomal (“RP”) and mitochondrial (“MT”) genes were next removed, yielding 3,147 highly variable genes which were subsequently analyzed using Seurat “RunPCA” function to obtain the first 20 principal components. Clustering was done using Seurat “FindClusters” function based on the 20 PCAs (resolution of 0.7). Visualization of the cells was performed using Uniform Manifold Approximation and Projection for Dimension Reduction (UMAP) algorithm as implemented by the Seurat “runUMAP” function and using the first 20 principal components.

#### Assigning cell type identity to single-cell clusters

Multiple methods were utilized to assign identities to derived cell clusters. The xCell (http://xcell.ucsf.edu/#)^71^ tool was used to compare the expression signatures of the initial 23 clusters with those of known cell types (n=64, including immune, endothelial, and stromal cells). Marker genes showing distinct expression in individual cell clusters compared to all others were identified using the Seurat FindAllMarkers function with default parameters. Marker genes with significant specificity to each cluster were compared to those reported elswhere^23, 24^.

#### Identification of cell-type maternal/fetal origin

We used two complementary approaches to determine the maternal/fetal origin of each cell-type. First, we used the samples derived from pregnancies where the neonate was male (3/9 cases, 8/25 samples) and we derived a fetal index based on the sum of all the reads mapping to genes on the Y chromosome relative to the total number of reads mapping to genes on the autosomes (Figure S4). The second method was based on the cell specific genotypes derived from the scRNA-seq read data and were overlapped to known genetic variants from the 1000 Genomes reference panel using the demuxlet and freemuxlet approach implemented in popscle^72^ (see Figure 1E), software available at https://github.com/statgen/popscle/.

#### Differential gene expression

To identify genes differentially expressed among locations (independent of study group), we created a pseudo-bulk aggregate of all the cells of the same cell-type. Only cell types with a minimum of 100 cell in each location were considered in this analysis. Differences in cell type specific expression were estimated using negative binomial models implemented in DESeq2^73^, including a fixed effect for each individual and location. The distribution of p-values for DEGs between pairs of compartments was assessed using a qq-plot to ensure the statistical models were well calibrated (Table S3). To detect DEGs across study groups we aggregated read counts across locations for each cell-type cluster, excluding cell-types with less than 100 cells in each study group (15 clusters). Differences in cell-type specific expression among study groups were estimated using negative binomial models implemented in Deseq2.

Differential gene expression was inferred based on FDR adjusted p-value (q-value <0.1) and fold change >2.0.

### Gene ontology and pathway enrichment analyses

The clusterProfiler^74^ package in R was utilized for the identification and visualization of enriched pathways among differentially expressed genes identified as described above. The functions “enrichGO”, “enrichKEGG”, and “enrichPathway” were used to identify over-represented pathways based on the Gene Ontology (GO), KEGG, and Reactome databases, respectively. Similar enrichment analyses were also conducted using Gene Set Enrichment Analysis (GSEA)^75^ which does not require selection of differentially expressed genes as a first step. Significance in all enrichment analyses were based on q<0.05.

### Quantification of single-cell signatures in maternal whole blood mRNA

#### Analysis of transcriptional signatures with advancing gestation and with labor at term

Whole-blood samples collected longitudinally (12 to 40 weeks of gestation) from women with a normal pregnancy who delivered at term with (TIL) (n=8) or without (TNL) (n=8) spontaneous labor, were profiled using DriverMap™ and RNA-Seq as previously described^58^. The log2 normalized read counts were averaged over the top genes (up to 20, ranked by decreasing fold change) distinguishing each cluster from all others as described above (single-cell signature). Whole blood single-cell signature expression in patients with three longitudinal samples was modeled using linear mixed-effects models with quadratic splines in order to assess the significance of changes with gestational age.

Differences in single-cell signature expression associated with labor at term (TIL vs. TNL) were assessed using two-tailed equal variance t-tests. In both analyses, adjustment for multiple signature testing was performed using the false discovery rate method, with q<0.1 being considered significant.

#### Analysis of transcriptional signatures in preterm labor

Whole blood gene expression data from samples collected at 24-34 weeks of gestation were profiled by RNA-Seq as previously described^59^. The study included samples from 15 women with preterm labor who delivered preterm, and 23 gestational age matched controls. Log2 transformed pseudo read count data were next transformed into Z-scores based on mean and standard deviation estimated in the control group. Single cell signatures were quantified as the average of Z-scores of member genes and compared between groups using a two-tailed Wilcoxon test. Adjustment for multiple signature testing was performed using the false discovery rate method, with q<0.1 being considered a significant result.

