## Supplementary Figures and Methods for "Single Cell Transcriptional Signatures of the Human Placenta in Term and Preterm Parturition"

### Supplementary Tables and Figures

Video S1: **3D reconstruction of the lymphatic endothelium in the decidua present in the CAM compartment**. Immunofluorescence co-expression of LYVE-1 (red) and CD31 (green) represents LED cells.

*See supplementary file:*
`file://VideoS1.mp4`
*also available at:*
`http://genome.grid.wayne.edu/LaborPlacenta/VideoS1.mp4`

Table S1: **Summary of the scRNA-seq libraries prepared.** Each row summarizes each 10X Genomics scRNA-seq library prepared and processed in this study: sample ID, number of cells detected after filtering, location of the tissue (BP = basal plate, PV = Placental Villi, CAM = chorioamniotic membranes), pregnancy condition (TNL = term no labor, TIL = term in labor, PTL = pre-term labor), gender of the neonate, and total number of UMIs detected.

| LibraryID | Total Cells | Location | Condition | Fetal Sex | Total UMI |
|---|---:|---|---|---|---:|
| s1DB | 2,574 | BP | TIL | Male | 15,122,851 |
| s1W | 2,984 | CAM | TIL | Male | 16,520,673 |
| s2DB | 2,313 | BP | TNL | Male | 7,897,722 |
| s2P | 2,490 | PV | TNL | Male | 13,393,737 |
| s2W | 2,905 | CAM | TNL | Male | 19,972,076 |
| s3DB | 2,340 | BP | TNL | Female | 9,158,823 |
| s3P | 2,546 | PV | TNL | Female | 21,869,861 |
| s3W | 2,038 | CAM | TNL | Female | 18,387,161 |
| s4DB | 3,165 | BP | TNL | Male | 19,265,820 |
| s4P | 3,007 | PV | TNL | Male | 24,326,111 |
| s4W | 2,691 | CAM | TNL | Male | 20,041,494 |
| s5DB | 4,629 | BP | TIL | Female | 16,519,752 |
| s5P | 3,012 | PV | TIL | Female | 14,238,071 |
| s5W | 2,011 | CAM | TIL | Female | 9,485,943 |
| s6DB | 3,723 | BP | PTL | Female | 25,713,932 |
| s6P | 1,238 | PV | PTL | Female | 12,637,713 |
| s6W | 4,826 | CAM | PTL | Female | 9,179,363 |
| s7DB | 2,945 | BP | PTL | Female | 24,058,875 |
| s7P | 2,711 | PV | PTL | Female | 27,913,872 |
| s7W | 3,221 | CAM | PTL | Female | 14,562,313 |
| s8DB | 3,054 | BP | PTL | Female | 17,932,348 |
| s8P | 4,246 | PV | PTL | Female | 46,082,098 |
| s8W | 5,232 | CAM | PTL | Female | 40,774,413 |
| s9DB | 3,992 | BP | TIL | Female | 20,688,976 |
| s9W | 4,013 | CAM | TIL | Female | 30,633,207 |

Table S2: **Summary of cell count by cell-type, location and condition.** Each row summarizes the total number of cells of each cell-type as determined by Seurat and split by pregnancy condition (TNL = term no labor, TIL = term in labor, PTL = pre-term labor), or location of the tissue (BP = basal plate, PV = Placental Villi, CAM = chorioamniotic membranes).

| | | Condition | | | Tissue | | |
|---|---|---|---|---|---|---|---|
| Cell-type | Total | TNL | TIL | PTL | BP | PV | CAM |
| B-cells | 1,176 | 435 | 265 | 476 | 450 | 408 | 318 |
| npiCTB | 2,405 | 888 | 282 | 1,235 | 660 | 1,740 | 5 |
| CTB | 11,743 | 4,139 | 1,310 | 6,294 | 3,489 | 8,127 | 127 |
| Decidual | 2,757 | 628 | 1,251 | 878 | 1,406 | 11 | 1,340 |
| Macrophage-1 | 3,761 | 842 | 1,281 | 1,638 | 1,633 | 376 | 1,752 |
| Macrophage-2 | 2,340 | 875 | 1,091 | 374 | 1,109 | 1,221 | 10 |
| Endometrial | 221 | 34 | 133 | 54 | 20 | 0 | 201 |
| Endothelial | 6,344 | 2,124 | 1,154 | 3,066 | 122 | 3 | 6,219 |
| EVT | 7,711 | 2,842 | 1,904 | 2,965 | 2,947 | 1,676 | 3,088 |
| Fibroblast | 669 | 120 | 240 | 309 | 405 | 129 | 135 |
| HSC | 509 | 99 | 169 | 241 | 206 | 63 | 240 |
| Monocyte | 2,668 | 258 | 1,114 | 1,296 | 1,236 | 979 | 453 |
| Stromal-3 | 636 | 0 | 585 | 51 | 0 | 0 | 636 |
| NK-cells | 8,124 | 2,413 | 2,027 | 3,684 | 3,746 | 935 | 3,443 |
| Stromal-1 | 1,764 | 443 | 710 | 611 | 36 | 3 | 1,725 |
| Stromal-2 | 2,474 | 1,514 | 650 | 310 | 1,129 | 1,344 | 1 |
| STB | 174 | 95 | 22 | 57 | 36 | 134 | 4 |
| T-cell-activated | 9,306 | 2,795 | 4,493 | 2,018 | 7,303 | 1,191 | 812 |
| T-cell-resting | 13,124 | 2,951 | 4,534 | 5,639 | 2,802 | 910 | 9,412 |

Table S3: **Marker Genes identified for each cell-type**. The columns represent: 1) Cluster or cell-type name, 2) Ensembl gene identifier, 3) Gene symbol, 4) pct.1 : percentage of cells in this cluster where the feature is detected, 5) pct.2 : percentage of cells in other clusters where the feature is detected, 6) log fold-change of the average expression between this cluster and the rest, 7) Nominal $p$-value, 8) Adjusted $p$-value (Bonferroni).
*See supplementary file:*
`file://TableS3.txt`
*also available at:*
`http://genome.grid.wayne.edu/LaborPlacenta/TableS3.txt`

Table S4: **Genes differentially expressed across compartments for each common cell-type**. The columns represent: 1) Cluster or cell-type name, 2) Comparison groups or contrast (i.e., BP vs PV, BP vs CAM, and CAM vs PV), 3) Ensembl gene identifier, 4) Gene symbol, 5) baseMean gene baseline expression as calculated by DESeq2, 6) $\log_2$ Fold Change of the first group in column 2 versus the second group, 7) Standard error estimated for the $\log_2$ Fold Change, 8) Nominal $p$-value, 9) $q$-value or adjusted $p$-value to control for FDR. Only rows with $q < 0.2$ are reported.
*See supplementary file:*
`file://TableS4.txt`
*also available at:*
`http://genome.grid.wayne.edu/LaborPlacenta/TableS4.txt`

Table S5: **Genes differentially expressed across conditions for each cell-type**. The columns represent: 1) Cluster or cell-type name, 2) Comparison groups or contrast (i.e., TNL vs TIL, TIL vs PTL), 3) Ensembl gene identifier, 4) Gene symbol, 5) baseMean gene baseline expression as calculated by DESeq2, 6) $\log_2$ Fold Change of the first group in column 2 versus the second group, 7) Standard error estimated for the $\log_2$ Fold Change, 8) Nominal $p$-value, 9) $q$-value or adjusted $p$-value to control for FDR. Only rows with $q < 0.2$ are reported.
*See supplementary file:*
`file://TableS5.txt`
*also available at:*
`http://genome.grid.wayne.edu/LaborPlacenta/TableS5.txt`

Table S6: **Summary of the sample demographics included in this study.** Data are given as medians with interquartile ranges (IQR) or as percentages (n/N). [a] One sample missing data.

| **Clinical Parameters** | **Term no labor (TNL)** | **Term in labor (TIL)** | **Preterm labor (TIL)** |
|---|---|---|---|
| Maternal age (years; median [IQR]) | 32 (28-35) | 25 (22-31.5) | 27 (23.5-29) |
| Body mass index (kg/m2; median [IQR]) | 31.2 (30.3-37.7) | 43.3 (36.9-44.1) | 32.5 (28.9-39.9) |
| Primiparity | 0% (0/3) | 66/7% (2/3) | 33.3% (1/3) |
| Cesarean section | 100% (3/3) | 66.7% (2/3) | 33.3% (1/3) |
| Gestational age at delivery (weeks; median [IQR]) | 39.6 (39.3-39.6) | 39.1 (38.8-39.9) | 35.1 (33.4-35.2) |
| Birthweight (g) | 4030 (3900-4160)[a] | 3310 (3160-3405) | 2145 (1680-2220.5) |
| **Ethnicity** | | | |
| African-American | 66.7% (2/3) | 66.7% (2/3) | 100% (3/3) |
| Caucasian | 33.3% (1/3) | 0% (0/3) | 0% (0/3) |
| Other | 0% (0/3) | 33.3% (1/3) | 0% (0/3) |

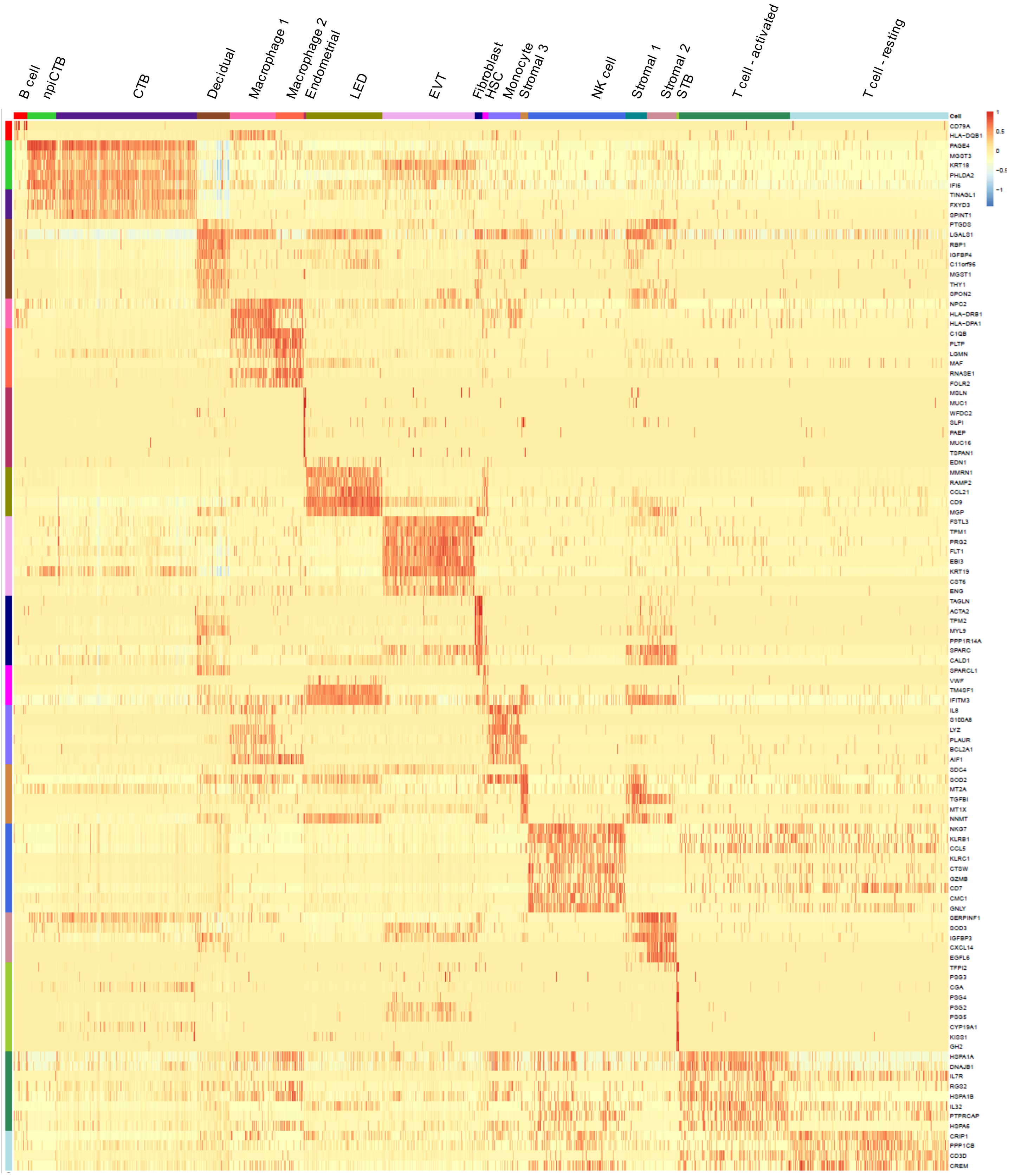


Figure S1: **Heatmap of the top gene expression markers defining each cell-type**. Each row represents a gene marker, and color represents normalized and scaled gene expression values derived from Seurat. Cell-type colors are consistent across all figures in this paper unless otherwise indicated.

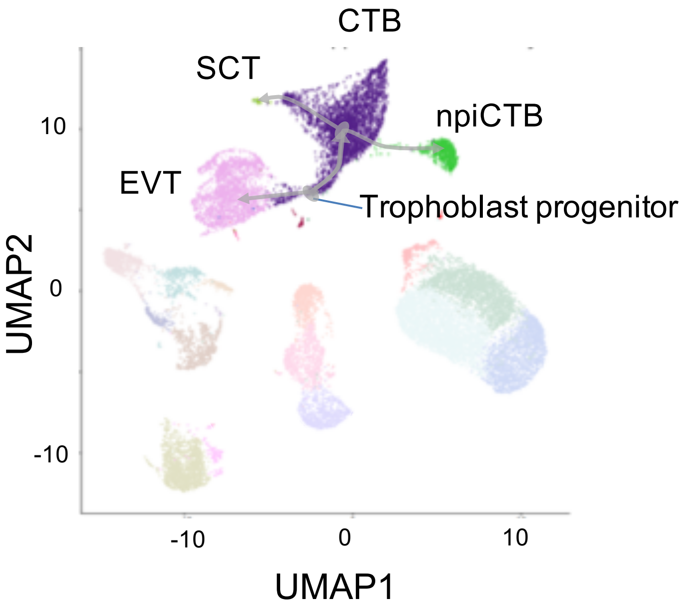


Figure S2: **UMAP plot highlighting the trophoblast cell-types and their inferred differentiation path**. Non trophoblast related cell-types are dimmed with a more saturated color. Lines and arrows reconstruct the most likely differentiation path of the different cell-types starting for cells that may be in a trophoblast progenitor state. The first branch seems to separate EVT from CTB, then CTB split in SCT and npiCTB.

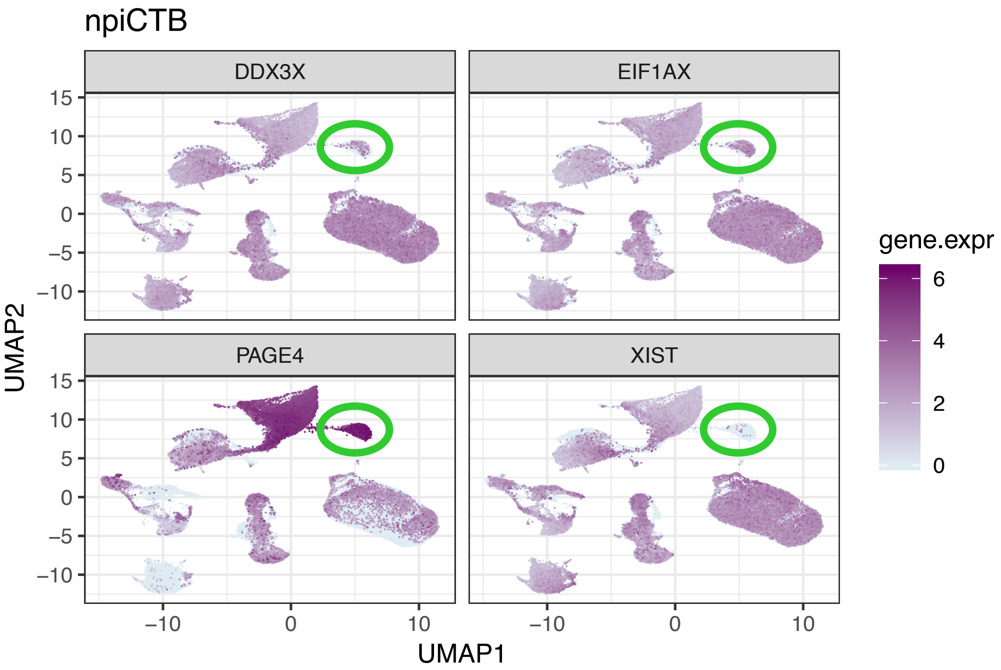


Figure S3: **Single marker gene expression UMAP plot for genes differentially expressed between CTB and npiCTB**. npiCTB cell-type is highlighted inside the green circle.

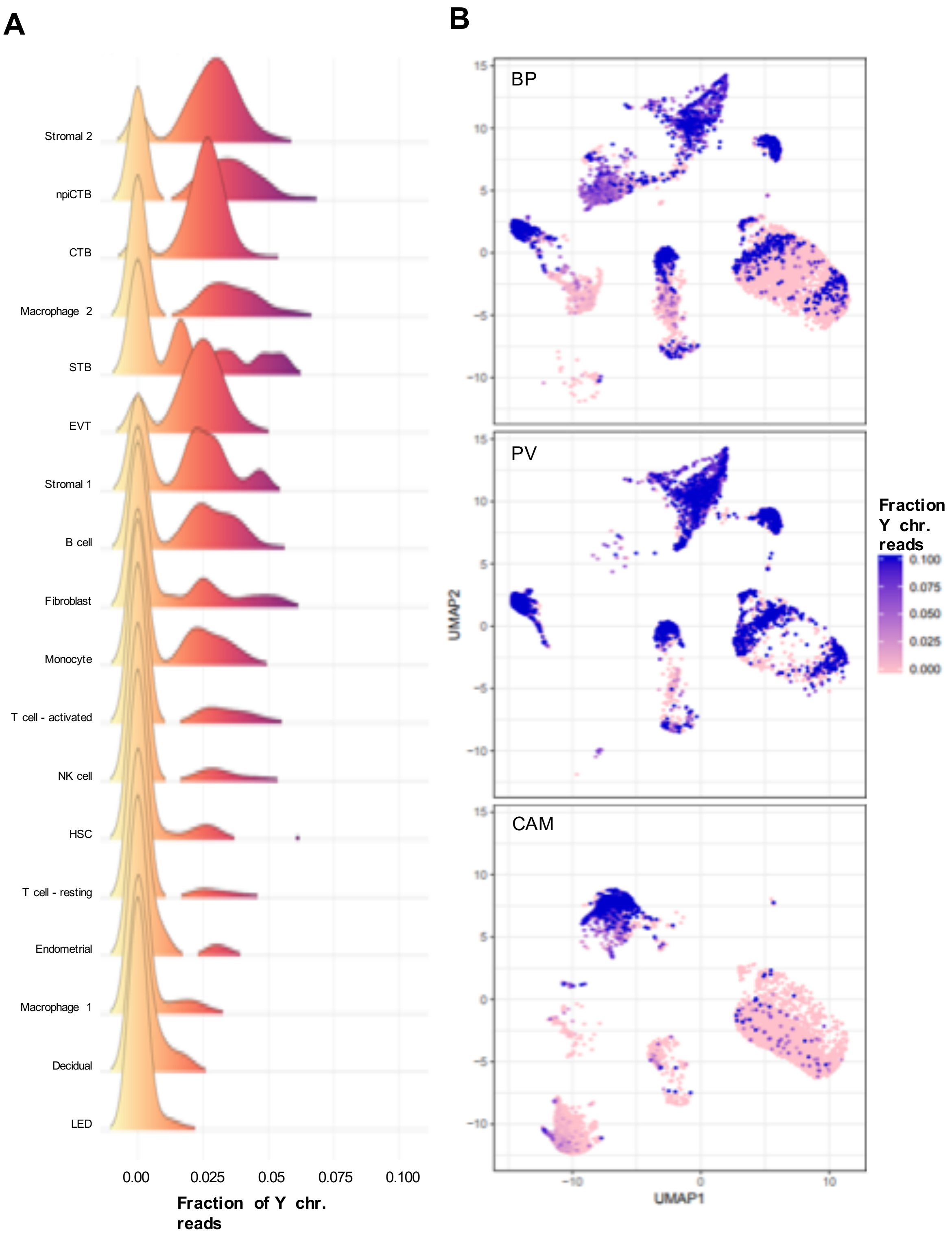


Figure S4: **Analysis of the Fetal/Maternal origin of the cell-types based on data from 3 pregnancies with a male fetus**. For each cell, an index is derived that represents the total number of reads mapping to the Y chromosome genes divided by the total number of reads mapping to autosomal chromosomes. (**A**) Density plot across all cells of the Y index. (B) UMAP plot where each facet represents a different placental compartment and each cell color is is scaled proportionally to the Y index.

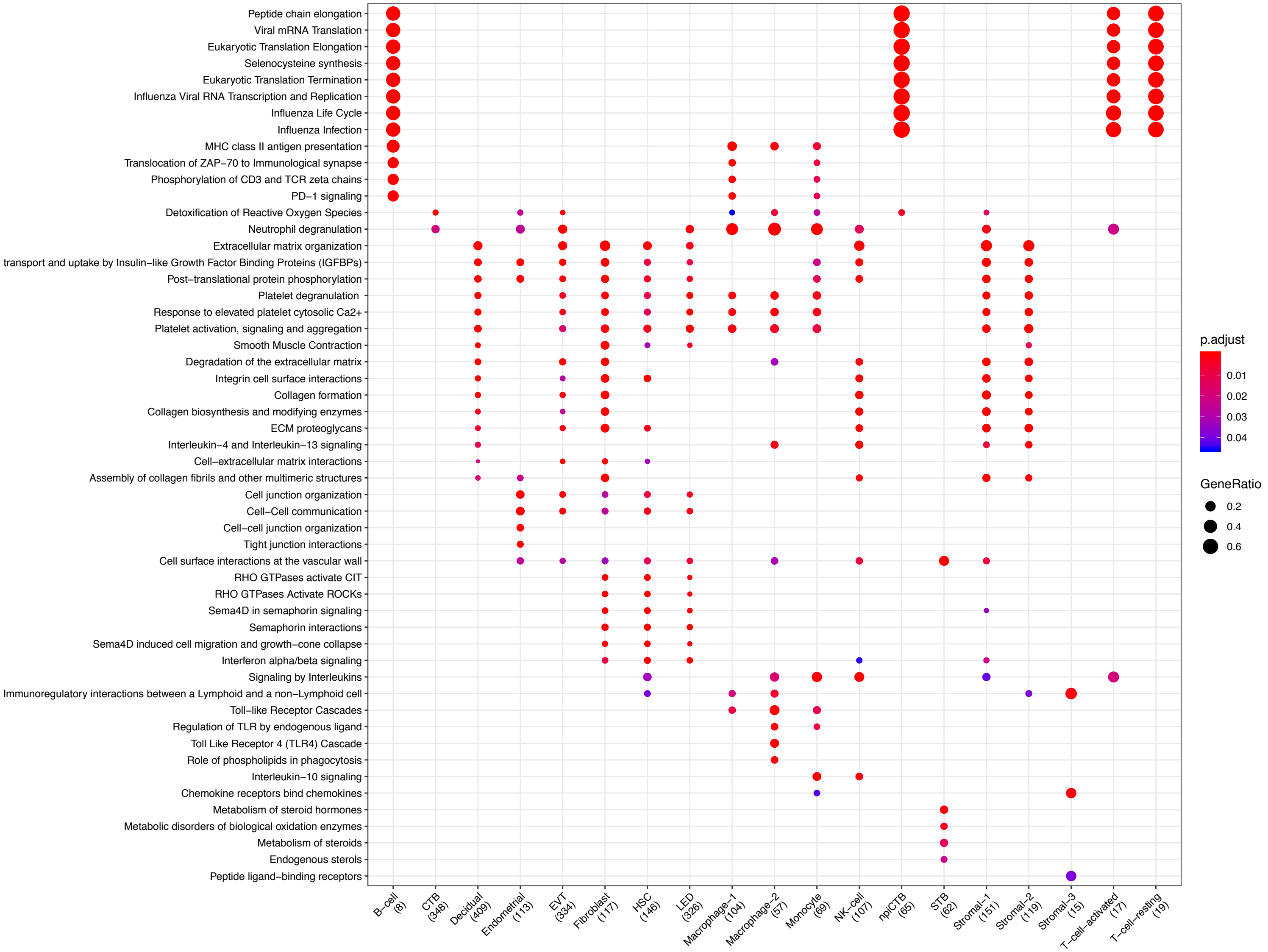


Figure S5: **Clusterprofiler dot plot showing the ReactomeDB Pathways enriched for genes that define each cell-type**. Color is scaled to the Benjamini Hochberg adjusted p-value, and dot size is scaled to the fraction of cell-type (column name) specific genes (number in parentheses) that are found annotated in the category (row name).

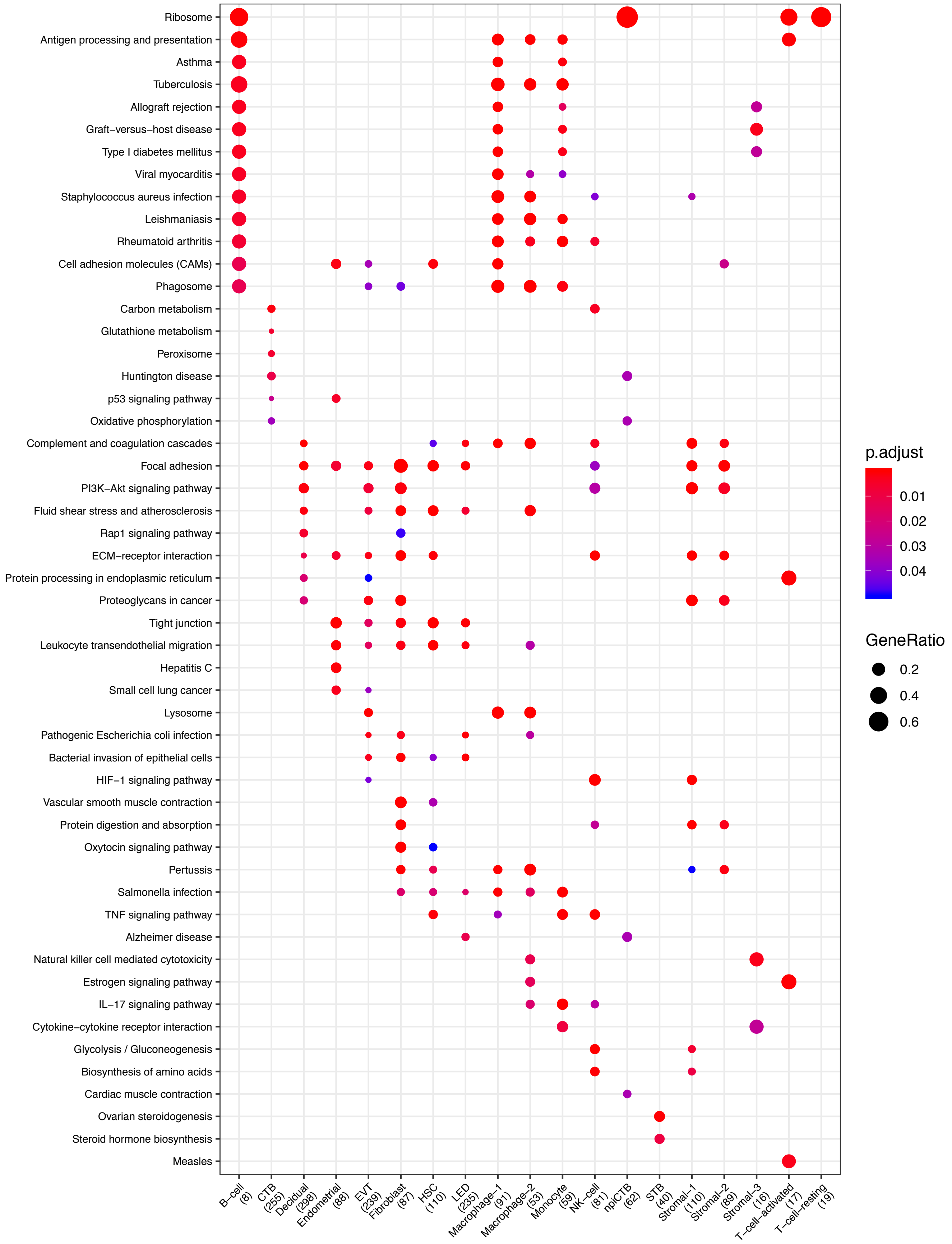


Figure S6: **Clusterprofiler dot plot showing the Kegg Pathways enriched for genes that define each cell-type**. Color is scaled to the Benjamini Hochberg adjusted p-value, and dot size is scaled to the fraction of cell-type (column name) specific genes (number in parentheses) that are found annotated in the category (row name).

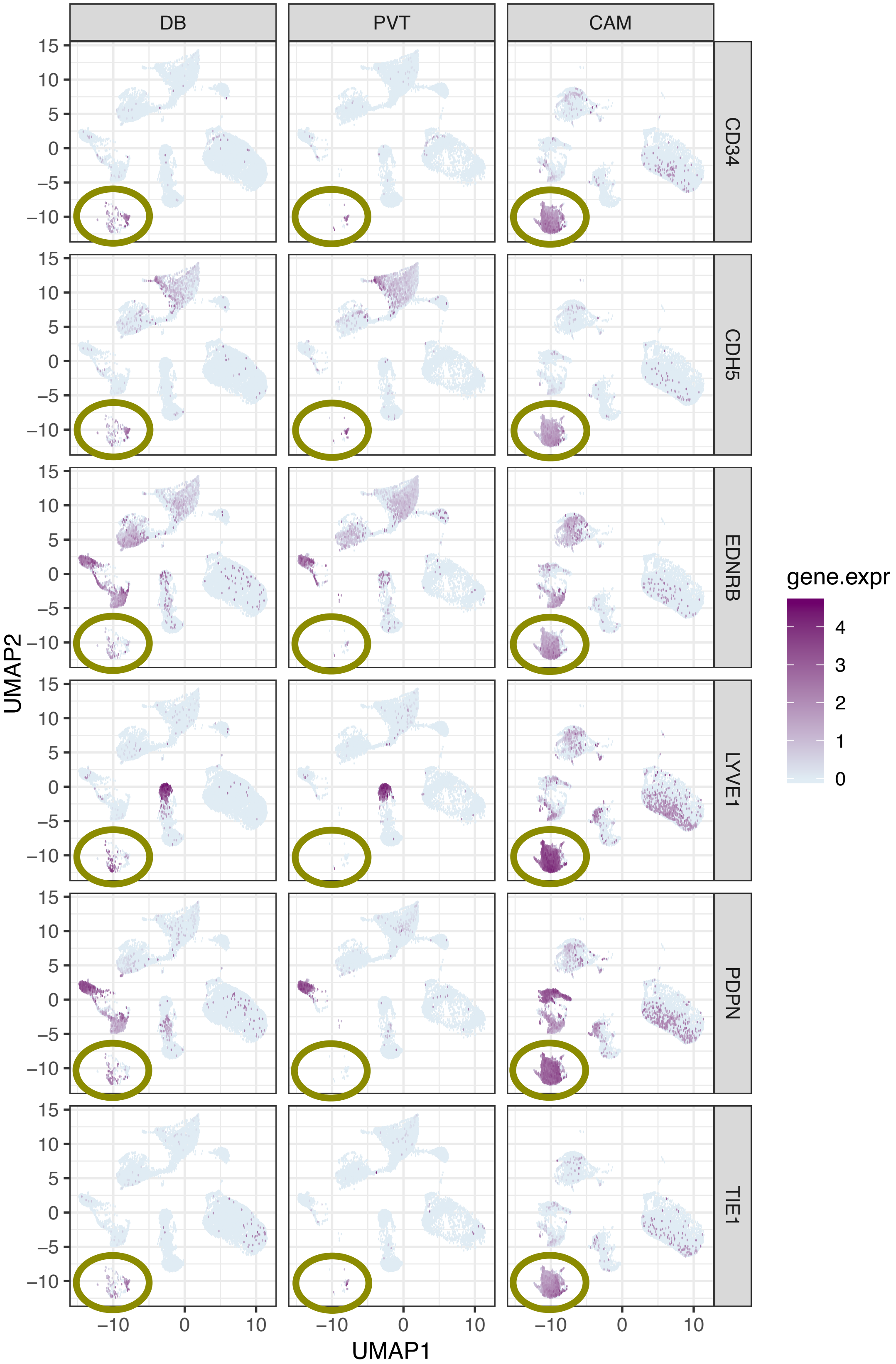


Figure S7: **Single marker gene expression UMAP plot for genes that are more highly expressed in lymphatic endothelial decidual (LED) cells**. Each row of panels represents a gene that is highly expressed in LEDs and column represents a compartments (Basal Plate = BP, Pacental Villi = PV, and chorioammiotic membranes = CAM). Note that LEDs highlighted inside the circle are almost only found in the CAM.

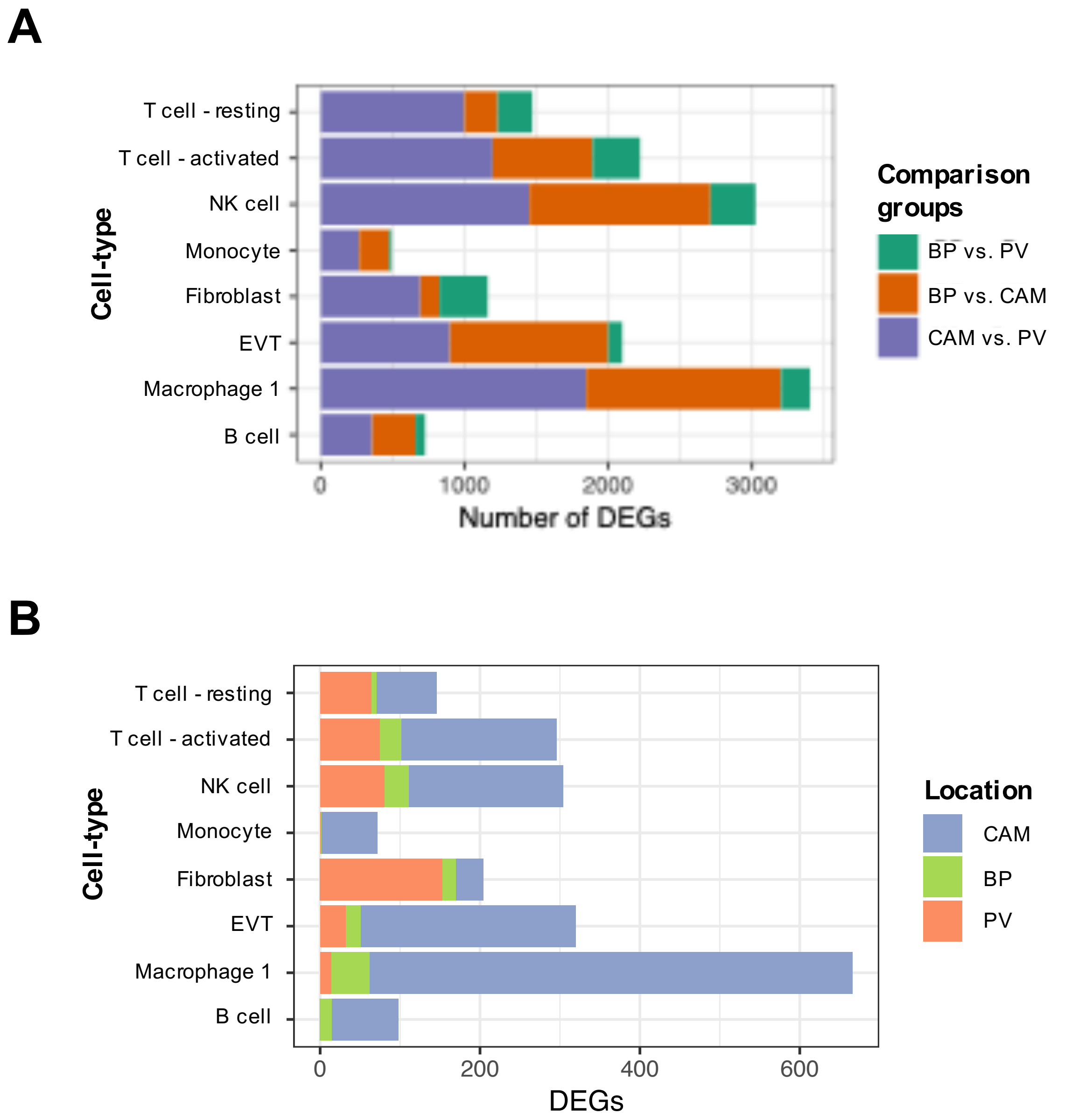


Figure S8: **Stacked bar plot summarizing differentially expressed genes across compartments for a cell types that are present on all three of them**. Number of DEGs at a Benjamini Hochberg adjusted p-value $< 0.1$ and fold change greater than 2: (**A**) between each pair of compartments, or (**B**) for each compartment (indicated in color) to the other two compartments.

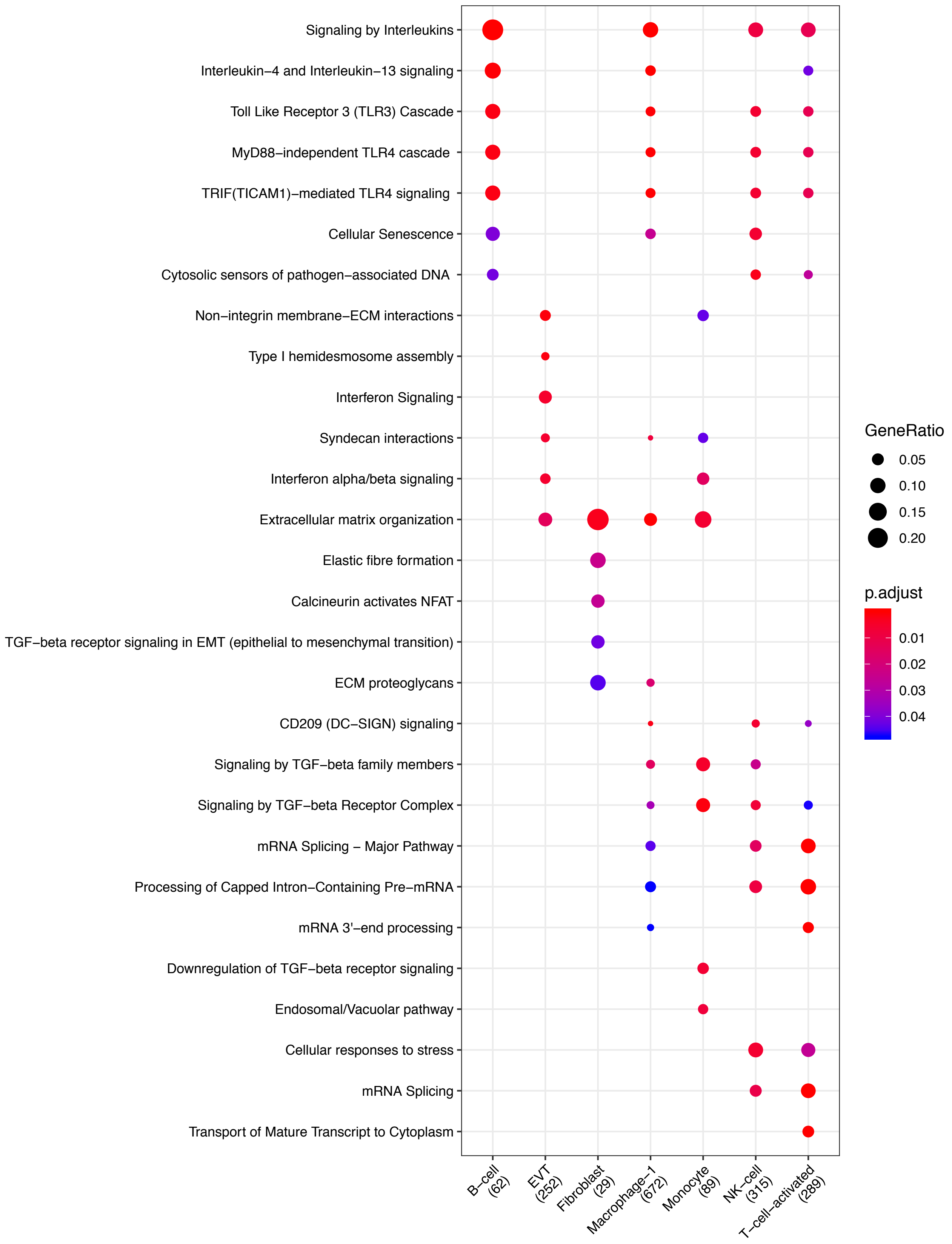


Figure S9: **Clusterprofiler dot plot showing the ReactomeDB Pathways enriched for genes that are significantly more highly expressed in the CAM compartment relative to the other compartments for each cell-type**. Color is scaled to the Benjamini Hochberg adjusted p-value, and dot size is scaled to the fraction of cell-type (column name) specific genes (number in parentheses) for the CAM that are found annotated in the category (row name).

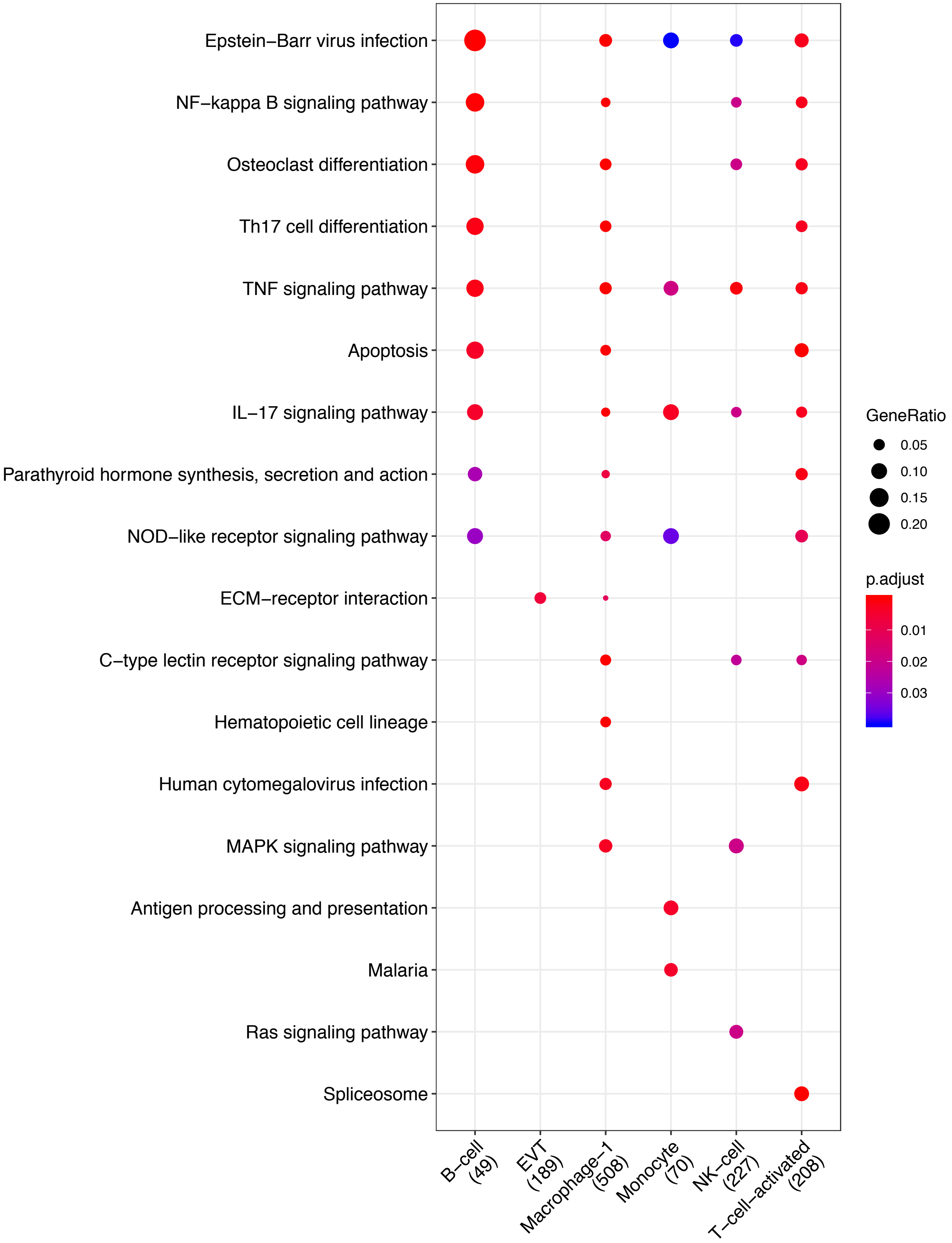


Figure S10: **Clusterprofiler dot plot showing the Kegg Pathways enriched for genes that are significantly more highly expressed in the CAM compartment relative to the other compartments for each cell-type**. Color is scaled to the Benjamini Hochberg adjusted p-value, and dot size is scaled to the fraction of cell-type (column name) specific genes (number in parentheses) for the CAM that are found annotated in the category (row name).

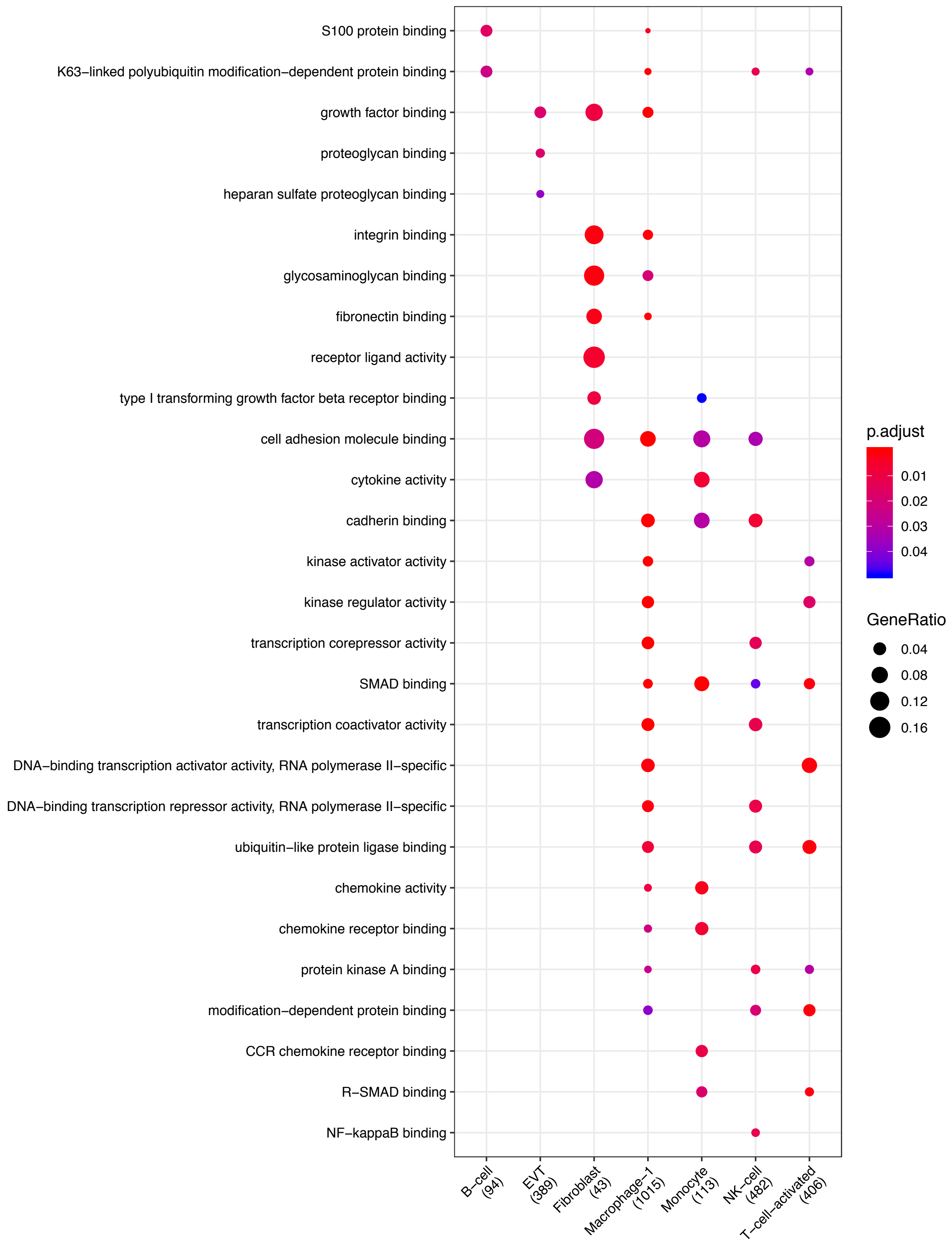


Figure S11: **Clusterprofiler dot plot showing gene ontology (GO) terms enriched for genes that are significantly more highly expressed in the CAM compartment relative to the other compartments for each cell-type**. Color is scaled to the Benjamini Hochberg adjusted p-value, and dot size is scaled to the fraction of cell-type (column name) specific genes (number in parentheses) for the CAM that are found annotated in the category (row name).

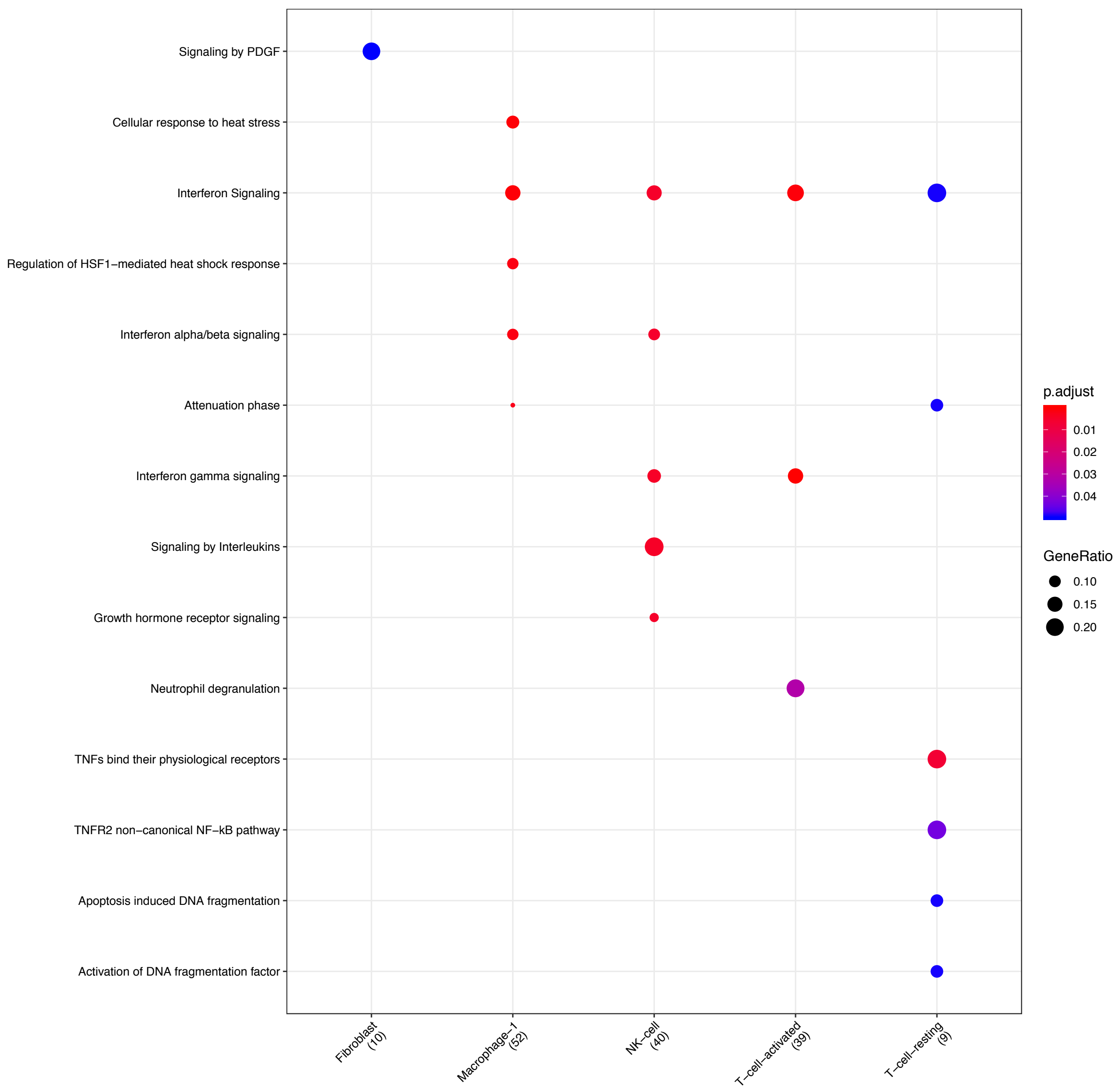


Figure S12: **Clusterprofiler dot plot showing the ReactomeDB Pathways enriched for genes that are significantly more highly expressed in the BP compartment relative to the other compartments for each cell-type**. Color is scaled to the Benjamini Hochberg adjusted p-value, and dot size is scaled to the fraction of cell-type (column name) specific genes (number in parentheses) for the BP that are found annotated in the category (row name).

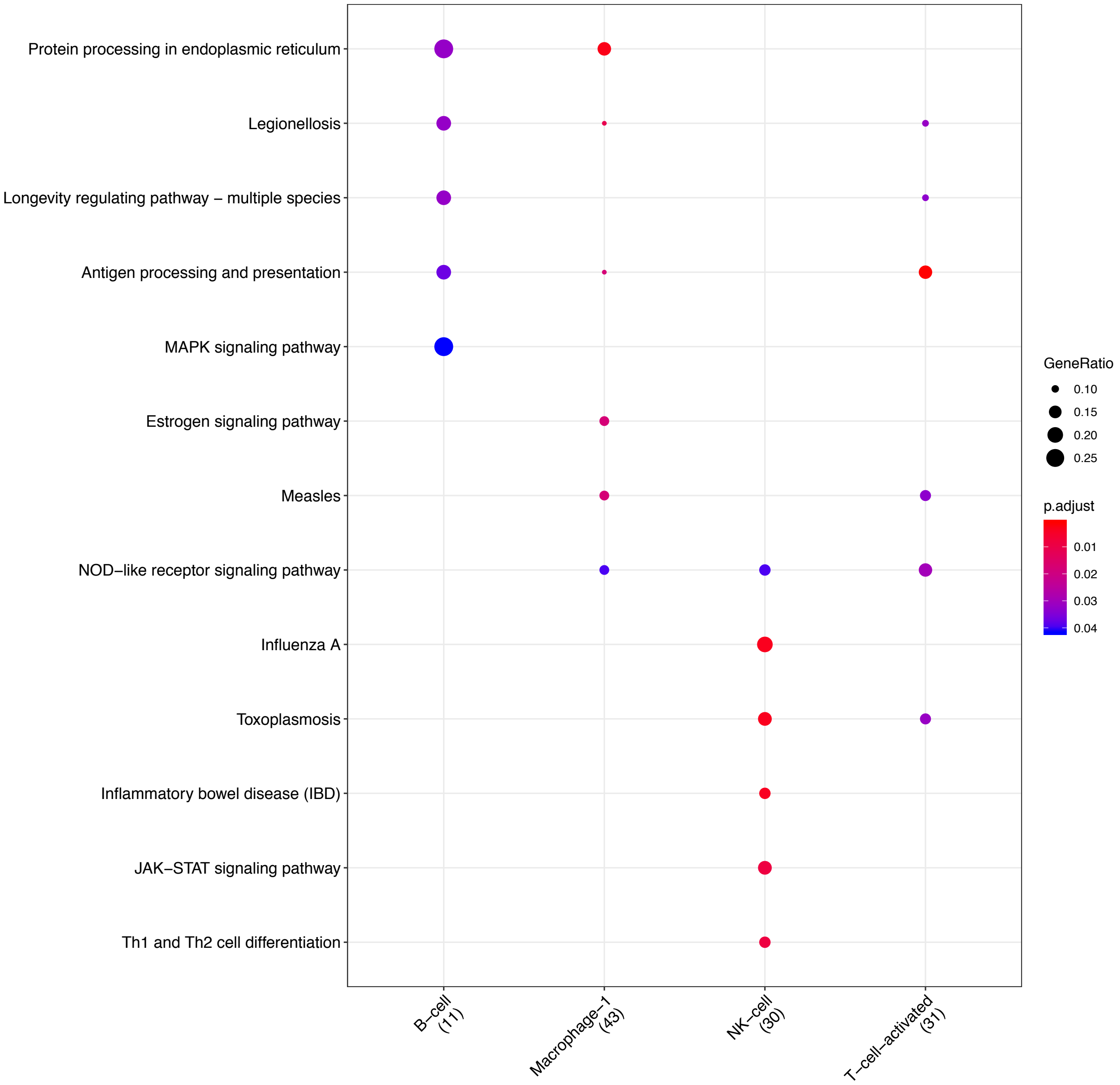


Figure S13: **Clusterprofiler dot plot showing the Kegg Pathways enriched for genes that are significantly more highly expressed in the BP compartment relative to the other compartments for each cell-type**. Color is scaled to the Benjamini Hochberg adjusted p-value, and dot size is scaled to the fraction of cell-type (column name) specific genes (number in parentheses) for the BP that are found annotated in the category (row name).

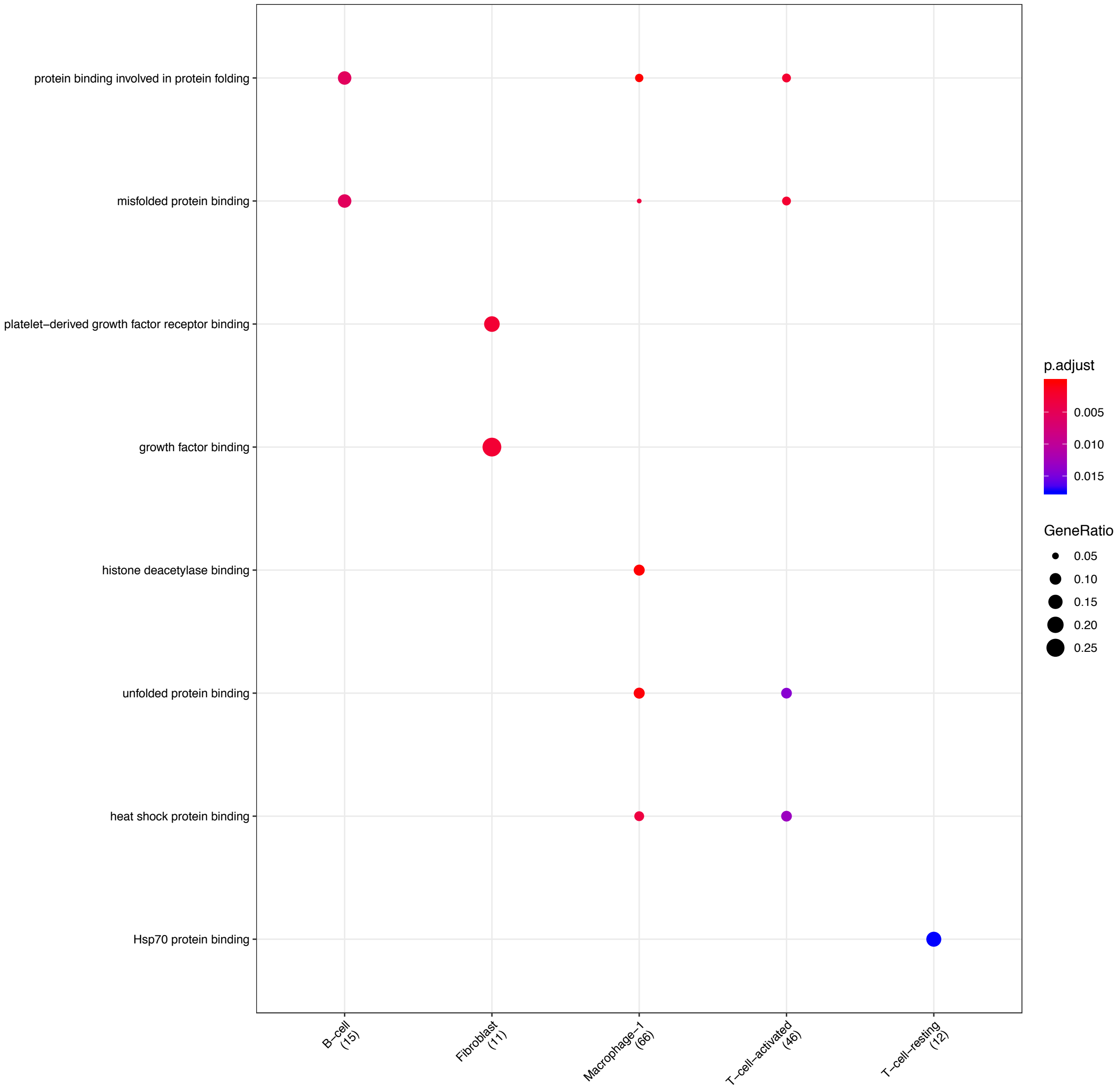


Figure S14: **Clusterprofiler dot plot showing gene ontology (GO) terms enriched for genes that are significantly more highly expressed in the BP compartment relative to the other compartments for each cell-type**. Color is scaled to the Benjamini Hochberg adjusted p-value, and dot size is scaled to the fraction of cell-type (column name) specific genes (number in parentheses) for the BP that are found annotated in the category (row name).

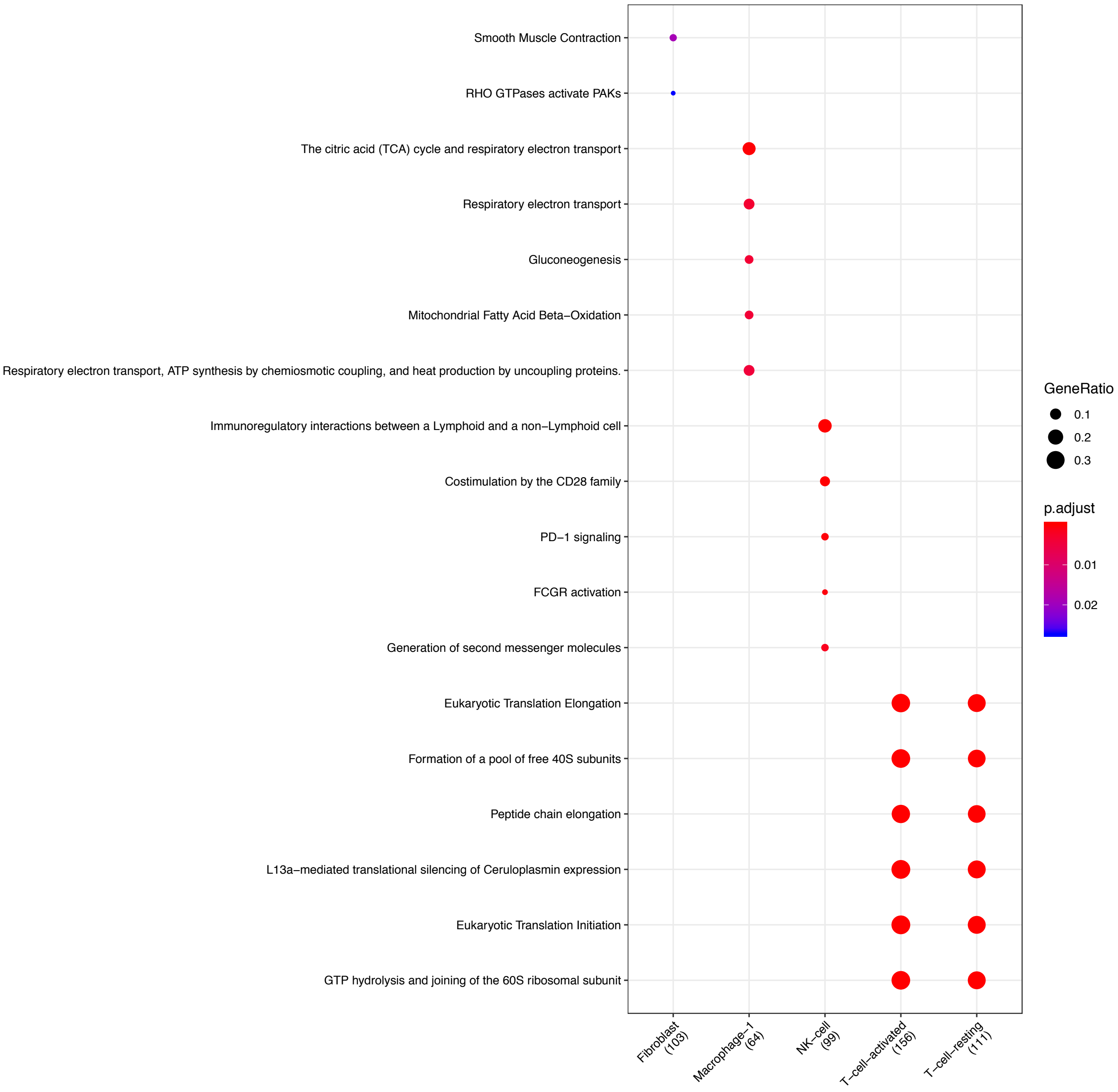


Figure S15: **Clusterprofiler dot plot showing the ReactomeDB Pathways enriched for genes that are significantly more highly expressed in the PV compartment relative to the other compartments for each cell-type**. Color is scaled to the Benjamini Hochberg adjusted p-value, and dot size is scaled to the fraction of cell-type (column name) specific genes (number in parentheses) for the PV that are found annotated in the category (row name).

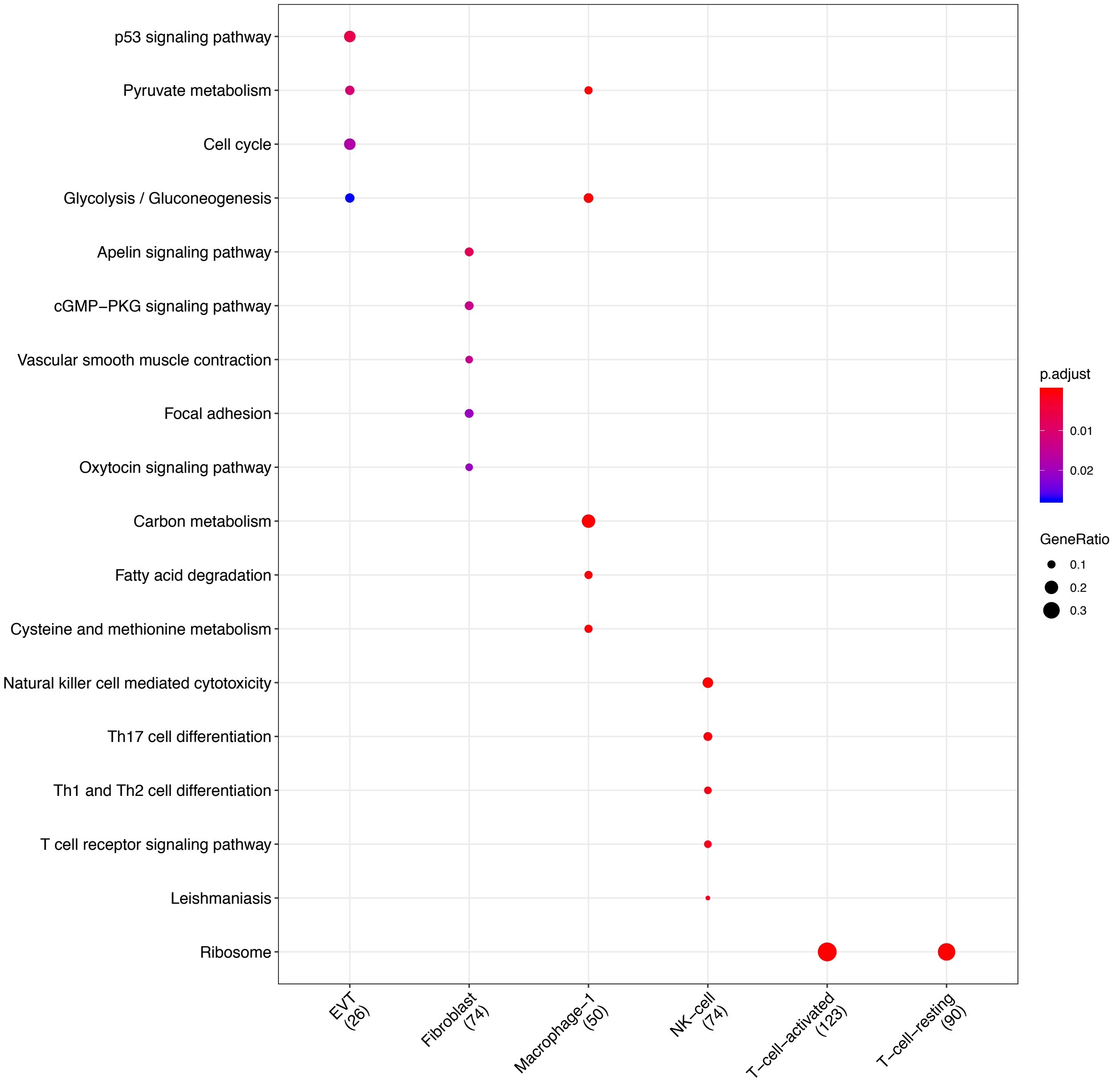


Figure S16: **Clusterprofiler dot plot showing the Kegg Pathways enriched for genes that are significantly more highly expressed in the PV compartment relative to the other compartments for each cell-type**. Color is scaled to the Benjamini Hochberg adjusted p-value, and dot size is scaled to the fraction of cell-type (column name) specific genes (number in parentheses) for the PV that are found annotated in the category (row name).

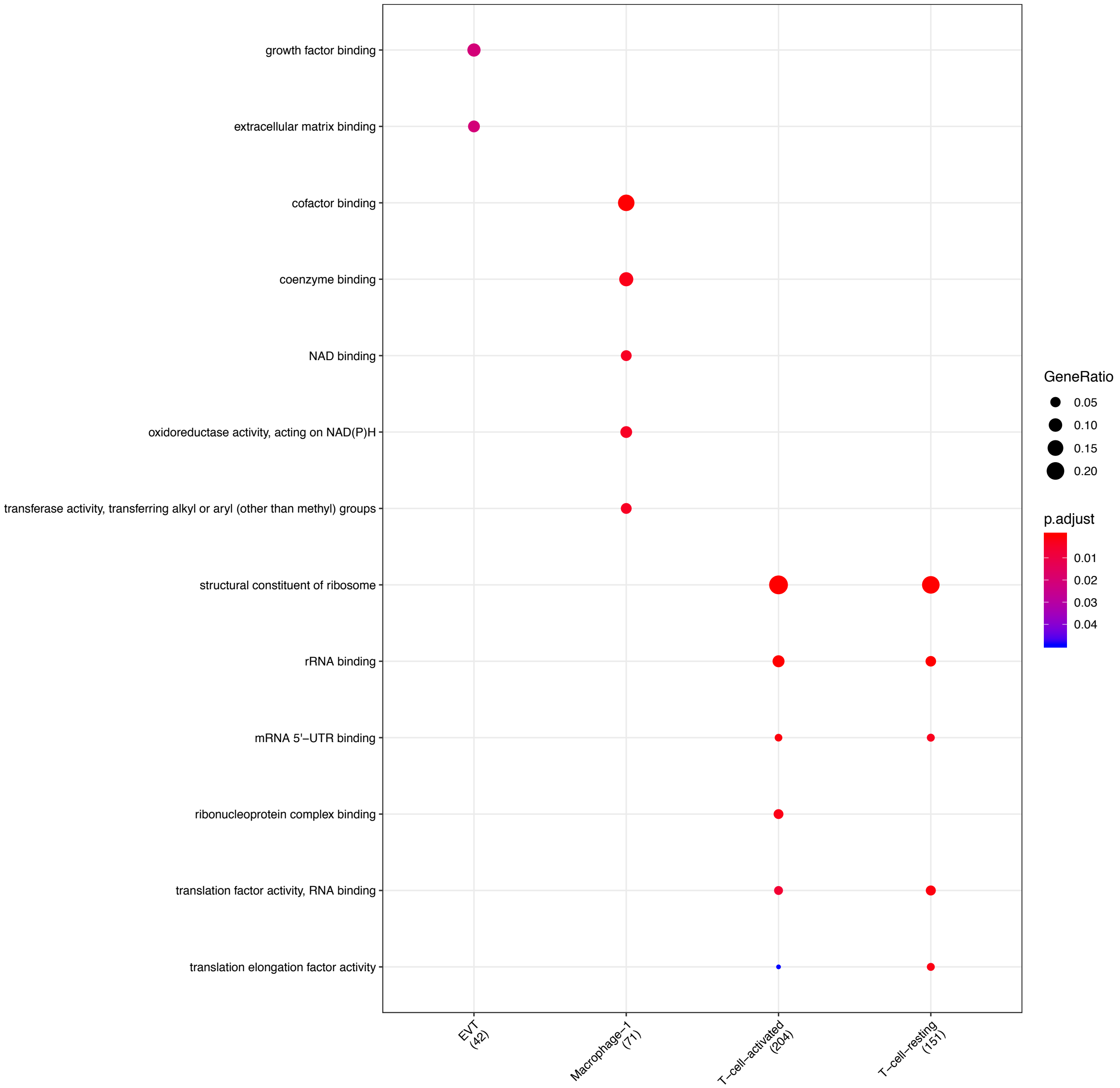


Figure S17: **Clusterprofiler dot plot showing gene ontology (GO) terms enriched for genes that are significantly more highly expressed in the PV compartment relative to the other compartments for each cell-type**. Color is scaled to the Benjamini Hochberg adjusted p-value, and dot size is scaled to the fraction of cell-type (column name) specific genes (number in parentheses) for the PV that are found annotated in the category (row name).

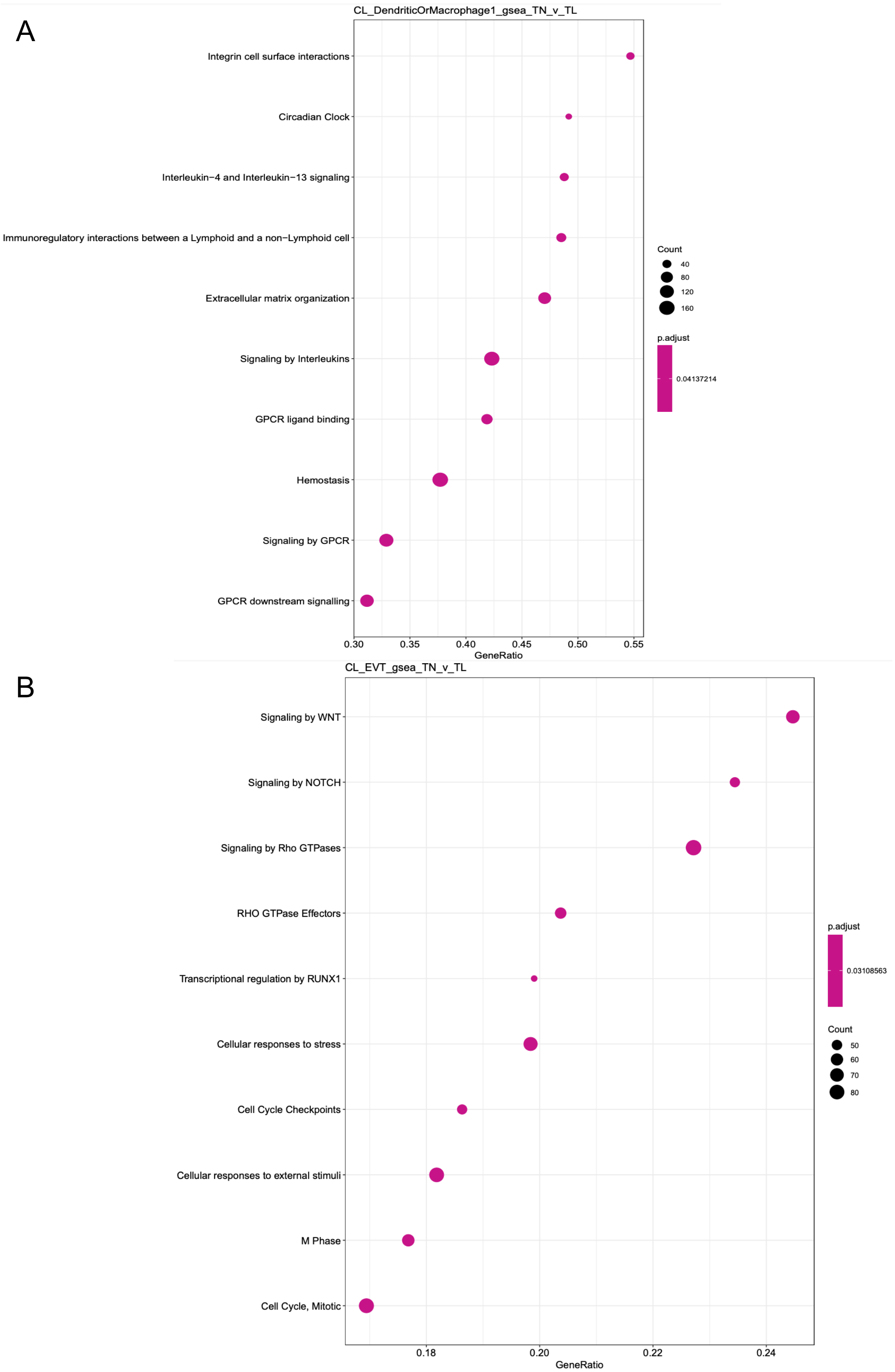


Figure S18: **Clusterprofiler dot plot showing ReactomeDB pathways enriched using gene set enrichment analysis (GSEA) for genes differentially expressed in term labor compared to term no labor condition**. The two panels correspond to the following cell-types: (**A**) maternal macrophages, and (**B**) extra villous trophoblasts (EVT).

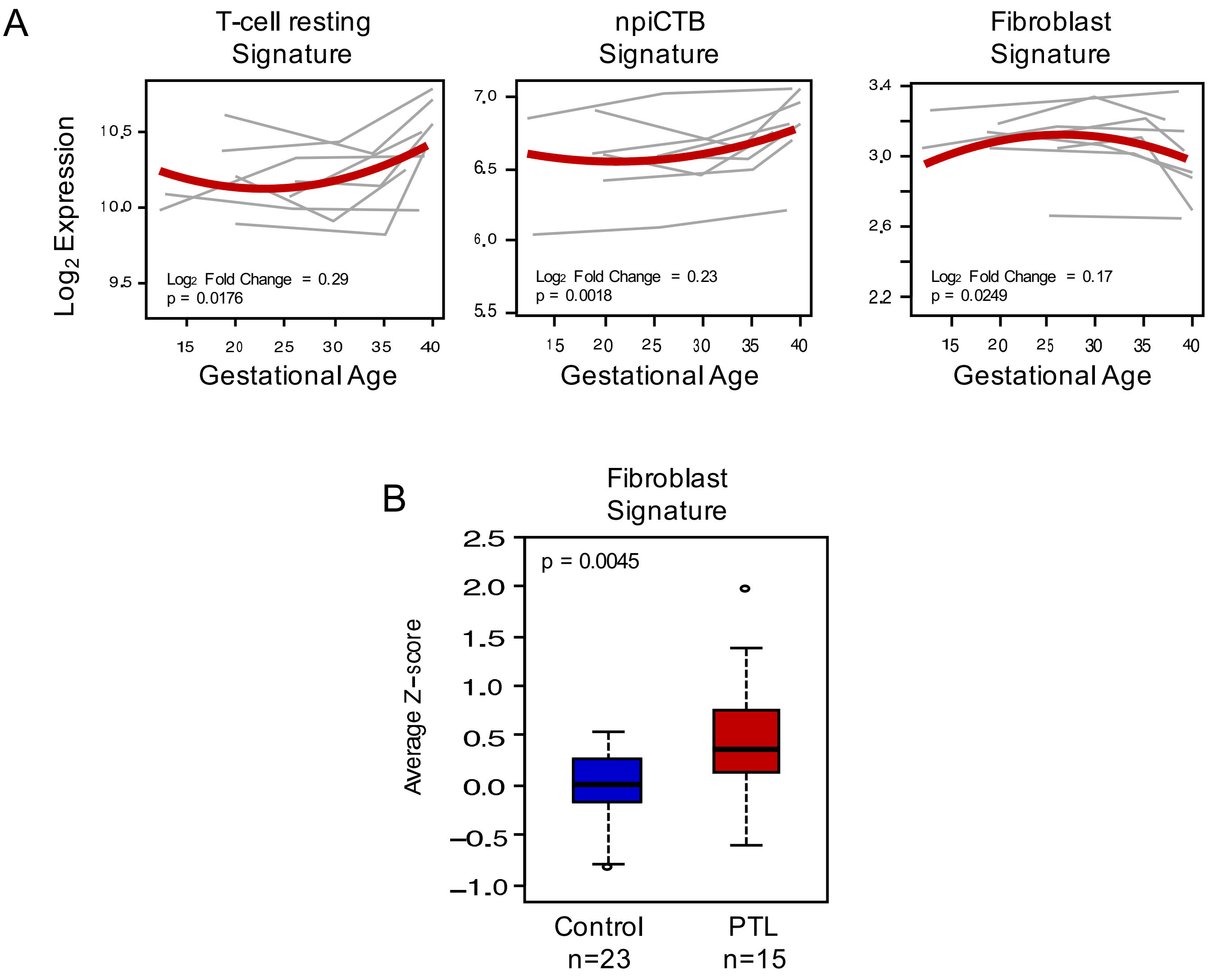


Figure S19: **Quantification of scRNA-seq signatures in maternal circulation**. Continued from main Figure 4 showing: (**A**) Expression of scRNA-seq signatures in the maternal circulation changing with advancing gestation; (**B**) perturbations in scRNA-seq signatures with preterm labor.
